# Neurophysiological and computational mechanisms of non-associative and associative memories during complex human behavior

**DOI:** 10.1101/2023.03.27.534384

**Authors:** Yuchen Xiao, Paula Sánchez López, Ruijie Wu, Ravi Srinivasan, Peng-Hu Wei, Yong-Zhi Shan, Daniel Weisholtz, Garth Rees Cosgrove, Joseph R Madsen, Scellig Stone, Guo-Guang Zhao, Gabriel Kreiman

## Abstract

The ability to transiently remember what happened where and when is a cornerstone of cognitive function. Forming and recalling working memories depends on detecting novelty, building associations to prior knowledge, and dynamically retrieving context-relevant information. Previous studies have scrutinized the neural machinery for individual components of recognition or associative memory under laboratory conditions, such as recalling elements from arbitrary lists of words or pictures. In this study, we implemented a well-known card-matching game that integrates multiple components of memory formation together in a naturalistic setting to investigate the dynamic neural processes underlying complex natural human memory. We recorded intracranial field potentials from 1,750 depth or subdural electrodes implanted in 20 patients with pharmacologically-intractable epilepsy while they were performing the task. We leveraged generalized linear models to simultaneously assess the relative contribution of neural responses to distinct task components. Neural activity in the gamma frequency band signaled novelty and graded degrees of familiarity, represented the strength and outcome of associative recall, and finally reflected visual feedback on a trial-by-trial basis. We introduce an attractor-based neural network model that provides a plausible first-order approximation to capture the behavioral and neurophysiological observations. The large-scale data and models enable dissociating and at the same time dynamically tracing the different cognitive components during fast, complex, and natural human memory behaviors.

## Introduction

Working memory serves as a fundamental component of our cognitive abilities, enabling us to store and retrieve immediate information. In stark contrast to most efforts in current artificial intelligence algorithms, the transient storage of memories occurs in a largely unsupervised fashion, with single or limited exposure. The formation and recall of memories require assessing novelty versus familiarity, building bridges between sensory inputs and prior knowledge, connecting spatial and temporal cues and effectively retrieving information in the context of current task demands. While substantial literature exists on neural responses in laboratory-based tasks for separate components of working memory, our understanding of how these components are integrated and coordinated in real-life tasks remains limited.

Non-associative recognition memory refers to the ability to judge the prior occurrence of a stimulus. Judging whether an item is novel or not is necessary for its successful memory encoding^1^, and recognizing an item as familiar facilitates memory retrieval^2^. Several studies have documented correlates of recognition memory for novelty versus familiarity, primarily but not exclusively, in medial temporal lobe (MTL) structures in rodents, monkeys, and humans^3–14^. Many studies have focused on tasks that involve presenting a list of items, such as words, pictures, or video clips, and either recalling items from these lists or assessing recognition memory for those items (e.g.,^6,8–12,15–20^). Both novel and familiar items need to be incorporated into the body of prior knowledge by forming novel associations. Associative memory refers to the ability to link items or evaluate the correctness of such associations (e.g.,^21–30^). Associative memory has been commonly investigated by having participants learn pairs of items and recalling one of the two items given the other or assessing whether a given association is correct or not.

Although recognition memory and associative memory have been largely studied separately, they are not independent in real-world memory tasks. The successful implementation of associative memory is contingent on basic recognition processes. To understand the connections and dissociations between different components of memory formation during natural and complex behavior, here we recorded intracranial field potentials from 20 patients with pharmacologically intractable epilepsy while they played a classical card matching game, colloquially known as the “Memory” game (**Figure 1, Movie S1**). We focused on the neural activity in the gamma frequency band (30-150 Hz)^31–35^. Participants thrived in the task, demonstrating dependence on the task memory demands and temporal recency effects. Using generalized linear models, we characterized how neural responses are modulated by the different behavioral components involved in the task. Our results demonstrate that neural circuits can represent novelty and familiarity independently of the sensory content, along with the strength and outcome of associative recall on a trial-by-trial basis.

**Figure 1.**
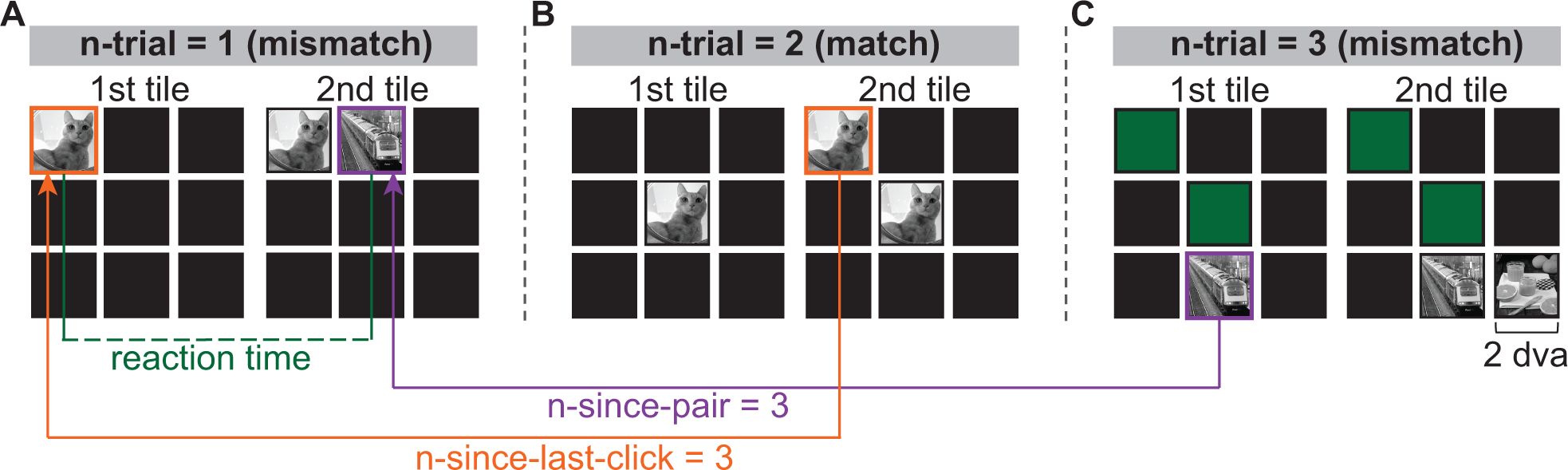
Experimental paradigm **A**-**C**. Three consecutive trials in a 3×3 board. In each trial, two tiles were flipped sequentially in a self-paced manner (1^st^ tile, then 2^nd^ tile). If the two tiles contained different images (**A**, **C**, **mismatch**), both tiles reset to their original active (black) state after 1 second. If both tiles contained the same image (**B**, **match**), they turned green after 1 second and stayed green for the remainder of the block. Three behavioral predictors used in the generalized linear models (GLM) are defined here: reaction time (the time between the 1^st^ and 2^nd^ tile within a trial), n-since-last-click (the number of clicks elapsed since the same tile was clicked last), and n-since-pair (the number of clicks elapsed since the last time a given tile’s matching pair was clicked). Each tile spanned approximately 2 degrees of visual angle (dva) in size. See also **Movie S1**.

To better understand the mechanisms underlying memory formation and retrieval, we turn to computational models^36^. Models rooted in persistent neuronal activity^37–45^ provide important insights into working memory’s neural basis. Recent perspectives, including those involving attractor networks^46–52^, have also highlighted the significance of Hebbian synaptic plasticity and short-term depression and facilitation as means to enhance memory encoding^53–57^. As a proof-of-principle, we introduce a simple attractor-based neural network model that provides a first-order approximation to describe the behavioral and neurophysiological observations.

## Results

We recorded intracranial field potentials (IFPs) from 20 patients with pharmacologically intractable epilepsy implanted with depth electrodes (**Table S1**, one participant also had subdural surface electrodes). Participants played a memory-matching game (**Figure 1, Movie S1, Methods**). Each trial consisted of two self-paced clicks. Clicking on a tile revealed an image (**Figure 1A**). Image categories included person, animal, food, vehicle, and indoor scenes. If the two tiles in a trial contained the same image (*match*, **Figure 1B**), the two tiles turned green and could not be clicked again for the remainder of the block. If the two images were different (*mismatch*, **Figure 1A** and **1C**), the two tiles turned black and could be clicked again. Participants started in a 3×3 tile board block like the one shown in **Figure 1** and progressed to more difficult blocks (4×4, 5×5, 6×6, or 7×7 tiles). All tiles had a corresponding match, except for one tile in the boards with an odd number of tiles (3×3, 5×5, and 7×7).

### Mismatch trials showed longer reaction times and were associated with less frequent and less recent exposure to matching pairs

The average number of clicks per tile increased with difficulty (board size), as expected (**Figure 2A**). All participants performed much better than a memoryless model (random clicking, p<0.001, here and in subsequent tests unless stated otherwise: permutation test, 5,000 iterations, one-tailed) and performed worse than a model assuming perfect memory (p<0.001, **Figure 2A**). The reaction time (RT) was defined as the time interval between the first and second clicks within a trial (**Figure 1A**). The reaction time was longer for mismatch than match trials for all board sizes (p<0.007, **Figure 2B**).

**Figure 2.**
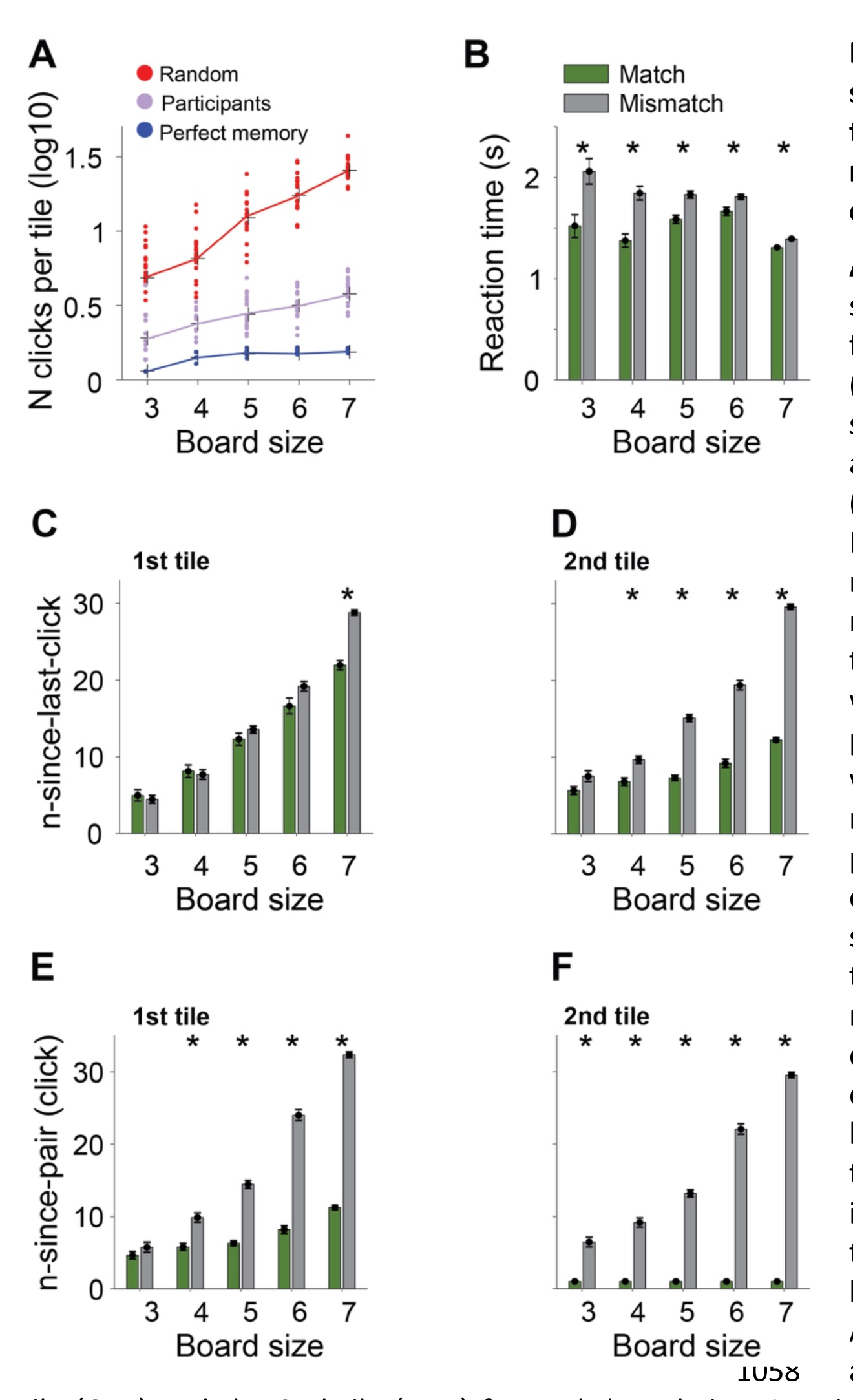
Behavioral measures show that participants thrived in the task and that finding matching pairs displayed classical memory effects **A.** Number of clicks per tile (log scale) as a function of board size for random simulation model (red, n=20), perfect memory simulation model (blue, n=20), and epilepsy patient participants (purple, n=20) (**Methods**). Perfect memory simulation models may generate different number of clicks per tile because the click location for new tiles was randomized. The performance of epilepsy patients was better than the random model and worse than the perfect model. The number of clicks per tile increased as board size incremented. **B.** Reaction times for match (green) and mismatch (gray) trials for different board sizes. Asterisks denote significant difference between match and mismatch trials (permutation test, 5,000 iterations, ⍺=0.01). Reaction time of mismatch trials was longer than match trials. **C-F.** Average n-since-last-click (**C**, **D**) and n-since-pair (**E**, **F**) for the 1st tile (**C**, **E**) and the 2nd tile (**D**, **F**) for each board size. Asterisks denote significant difference between match and mismatch trials (permutation test, 5,000 iterations, ⍺=0.01). For n-since-last- click, trials in which a tile was clicked for the 1st time were excluded in this figure. For n-since- pair, trials in which any tile’s matching pair had not been seen before were excluded in this figure. All error bars indicate s.e.m. (n=20 participants).

For a tile in a given trial, we defined *n-since-last-click* (nslc) as the number of clicks elapsed since the last time *the same tile* was clicked (**Figure 1A-B**). As expected, nslc increased with board size (p<0.001, linear regression, F-test, **Figure 2C-D**). For the 2^nd^ tile, nslc was larger in mismatch compared to match trials for all board sizes except the 3×3 case (p<0.001, **Figure 2D**), a reflection of the decay in memory for tiles that were not seen recently. The larger nslc in mismatch trials also held for the 1^st^ tile only for the 7×7 board size (p<0.001, **Figure 2C**). If the participants believe that they know the locations of both tiles in a matching pair, a reasonable strategy is to click first the tile they are less sure about, likely because they have seen this tile earlier rather than later in the block. This strategy accounts for the differences between the 1^st^ tile (**Figure 2C**) and 2^nd^ tile (**Figure 2D**).

For a tile in a given trial, we defined *n-since-pair* (nsp) as the number of clicks since the last time when *its matching pair* was seen (**Figure 1A**, **C**). As expected, nsp increased with board size given the increased difficulty (p<0.001, linear regression, F-test, **Figure 2E-2F**). Additionally, the more recent the tile’s matching pair was seen, the more likely the trial was a match. Thus, nsp was larger in mismatch compared to match trials in all cases except the 3x3 board size for the 1^st^ tile (p<0.001, **Figure 2E**). For the 2^nd^ tile, nsp for any match trial was always one because the matching pair would have been revealed in the previous click, by definition. Thus, there was a large difference between nsp between match and mismatch trials (p<0.001, **Figure 2F**).

We performed infrared eye-tracking on ten healthy participants while they performed the same task. Participants fixated on the tile they clicked, both for the first and second tiles, and both for match and mismatch trials (**Figure S1**). For the first tile, there was no difference in the dynamics of saccades towards and away from the target tile between match and mismatch trials (**Figure S1A**) after equalizing the RT and the distances between the 1^st^ and the 2^nd^ tiles. For the second tile, there was no difference in the dynamics of saccades toward the target tile before the click. However, within the 1 second window after the 2^nd^ click, the distance to the center of the tile was, on average, 1.76 dva (degrees of visual angle) larger for match than mismatch trials (**Figure S1B**). This small difference may be attributed to participants’ lingering slightly longer during mismatch trials, arguably in an effort to remember the tile.

### Neural signals reflect novelty and familiarity

We recorded intracranial field potentials from 1,750 electrodes (**Table S2**). We excluded 582 electrodes due to bipolar referencing, locations in pathological sites, or signals containing artifacts (**Methods**). We included in the analyses 676 bipolarly referenced electrodes in the gray matter (**Figure S2**) and 492 in the white matter (**Figure S3**). **Table S2** describes electrode locations separated by brain region and hemisphere. Although the white matter is presumed to contain mostly myelinated axons, previous studies have shown that intracranial field potential signals from the white matter can demonstrate biologically meaningful information^58,59^. Such signals could reflect small errors in electrode localization on the order of ∼2 millimeters and also the spread of intracranial field potential signals over 1 to 5 millimeters^32,33,60–62^, implying that white matter electrodes may still capture activity from gray matter. Indeed, we show here that electrodes in the white matter reveal task-relevant properties and therefore included electrodes in the white matter in our analyses. To avoid confusion about the origin of the signals, we focused on the gray matter electrodes in the main text and reported results from electrodes in the white matter in the Supplementary Material. None of the conclusions in this study would change if we were to report the results from electrodes in the gray matter exclusively.

We built two generalized linear models (GLM) to characterize how the neural responses depended on the cognitive demands of each trial. The first model focused on the neural responses to the 1^st^ tile, and the second model on the 2^nd^ tile. In both cases, we focused on predicting the area under the curve (AUC) of the gamma band power (30-150 Hz) in each trial (**Methods**). For the first GLM, the time window started when the 1^st^ tile was clicked and ended at a time corresponding to the 90th percentile of the distribution of reaction times (time difference between the 1^st^ and the 2^nd^ clicks, **Figure 1A**). This criterion was a reasonable tradeoff between minimizing overlap with responses after the 2^nd^ tile and maximally capturing information before the 2^nd^ tile. For the second model, the time window started with the 2^nd^ click and ended one second afterward.

We considered 15 predictors for the GLM models, including whether a trial was a match or not, reaction time, n-since-last-click (nslc), and n-since-pair (nsp), the variables introduced in **Figures 1**-**2**. We also included additional predictors: first-click (whether a tile was clicked for the first time), n-times-seen (number of times an image had been seen), next-match (whether the subsequent trial was a match), board size, x and y position of the clicked tile within the board, distance between the first and the second tiles, and whether the image contained a person, animal, food, or vehicle. **Table 1** lists all the predictors and their definitions. Since several predictors were correlated with each other (**Figure S4A**-**B**), we computed the variance inflation factor (VIF)^63^, a metric commonly used to account for correlations between predictors in generalized linear models. The VIF of each predictor was smaller than 3 for all participants (**Figure S4C**-**D**). Therefore, the correlations between predictors did not harm the performance of the models (**Methods**).

**Table 1.**
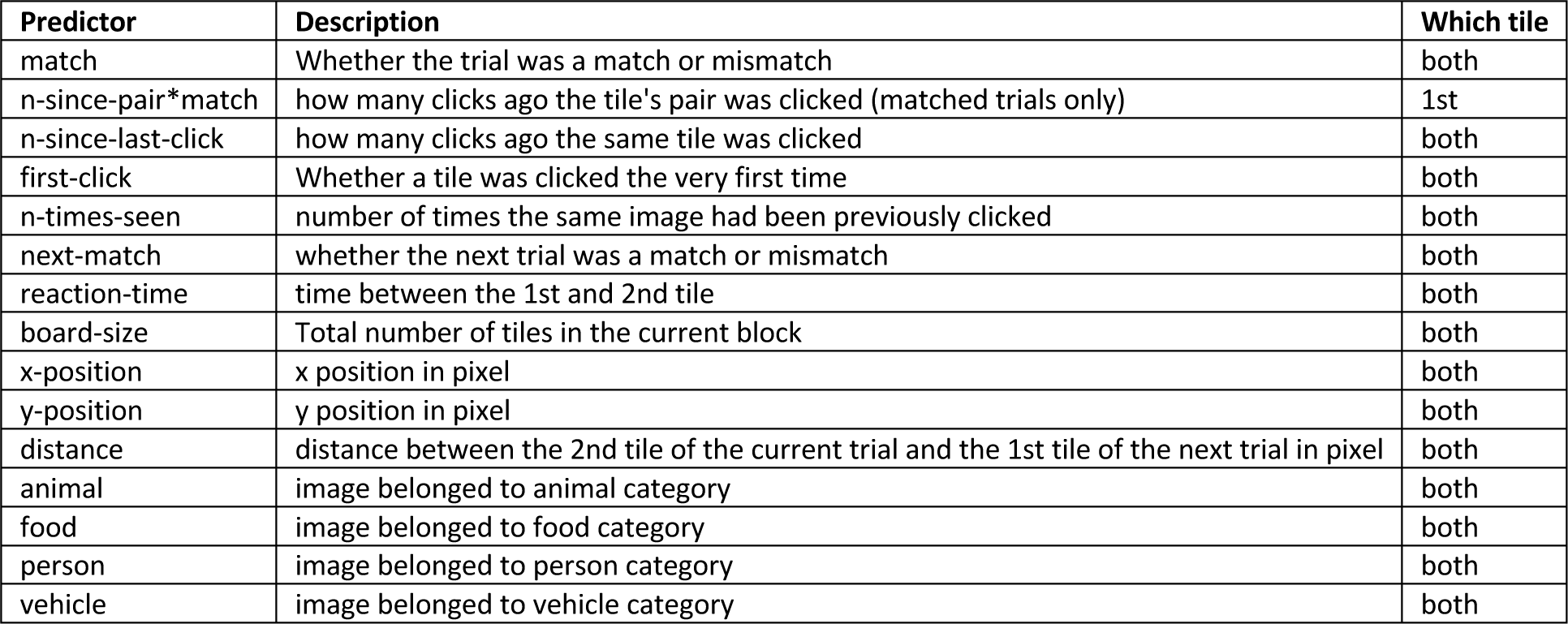
Predictors in the generalized linear models, their definitions, and applicable tiles.

When the first tile in a trial was clicked, its status in memory guided the following actions. If it was a new image, the participant needed to encode it in memory for future retrieval. Thus, the ability to detect novelty is the first step for successful encoding. The predictor *first-click* described novelty and had a value of 1 whenever a tile was seen for the first time and 0 otherwise. If a tile had been viewed before, it would appear familiar to the participant, and the degree of familiarity depended on how long ago that tile had been seen last. The predictor n-since-last-click (**Figure 1A-B**, **Figure 2C-D**) captures the notion of familiarity; the smaller the nslc value, the more familiar the tile is because that same tile was seen more recently and there were fewer competing stimuli encountered in between.

Figure 3A-D shows the neural activity of an electrode located in the right lateral orbitofrontal cortex (arrow in Figure 3D), whose responses to the first tile correlated with novelty. The GLM analysis indicated that first-click was a significant predictor of the neural responses (Figure 3A). The neural responses to the first tile showed a decrease in activity for novel tiles compared to tiles that had been seen before (Figure 3B), which could also be readily seen in individual trials (Figure 3C). This decrease in activity is reflected by the negative sign in the GLM first-click predictor (Figure 3A).

**Figure 3.**
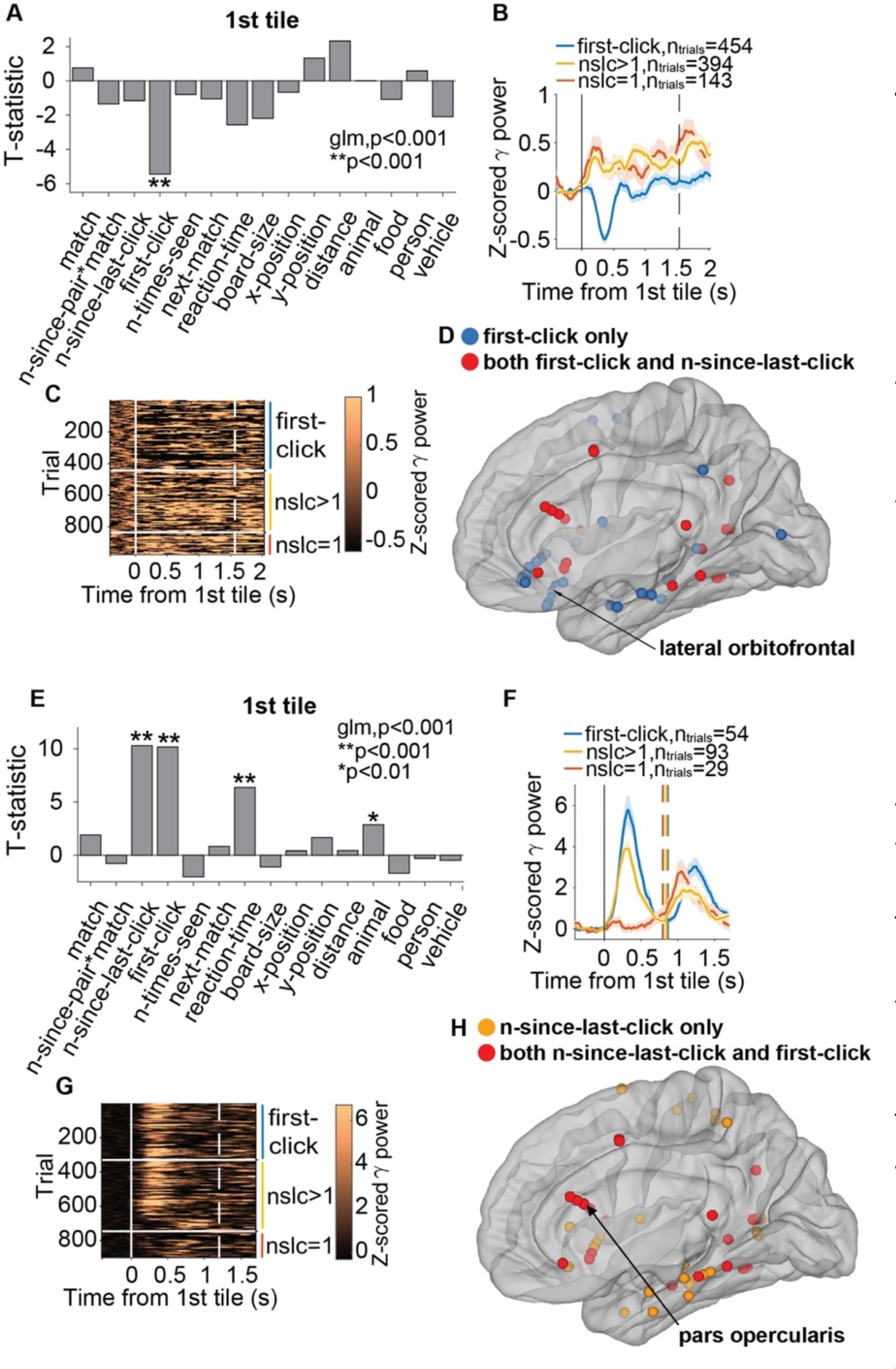
Neural signals reflect novelty and familiarity Panels show two example electrodes, one in the right lateral orbitofrontal cortex (**A**-**D**), one in the left pars opercularis (**D-H**), and population locations in **D**, **H**. **A**, **E**. T-statistic of each predictor in the GLM analyses (**Methods**). Asterisks indicate statistically significant predictors for the neural signals. **B**, **F**. Z-scored gamma band power aligned to the 1^st^ tile onset (solid vertical line) for novel tiles (blue), unfamiliar tiles (n-since-last-click>1, yellow), and familiar tiles (n-since-last-click=1, red). The vertical dashed line indicates the mean reaction time. Multiple dashed lines in **F** indicate reaction time equalization (**Methods**). The time axis extends from 400 ms before the click to 500 ms after the average reaction time. **F** displays only trials after RT equalization (**Methods**). Shaded error bars indicate s.e.m. **C, G.** Raster plots showing the z-scored gamma power in individual trials ordered by first-click and then larger to smaller n- since-last-click; division indicated by white horizontal lines/spaces and colored vertical bars. **D.** Locations of all electrodes where first-click was a significant predictor during the 1st tile. Blue: first-click only; red: both first-click and n-since-last-click were significant predictors. **H.** Locations of all electrodes where n-since-last-click was a significant predictor during the 1st tile. Orange: n-since-last-click only; red: both n-since-last-click and first-click were significant predictors. All electrodes were reflected on one hemisphere for display purposes.

Novelty was a significant predictor of the neural responses after the first tile (p<0.01, GLM) for 50 electrodes in the gray matter (7.4% of the total, **Table S3A**, Figure 3D) and 33 electrodes in the white matter (6.7% of the total, **Table S3B**). The lateral orbitofrontal (LOF) cortex and pars opercularis contained significantly more electrodes than expected by chance (p<0.01, **Methods**).

Figure 3E-H shows the neural activity of an electrode located in the left pars opercularis (arrow in Figure 3H), whose responses to the first tile correlated with familiarity. The GLM analysis indicated that both n-since-last-click (nslc) and first-click were significant predictors of the neural responses (p<0.001, GLM, Figure 3E). Novel tiles (completely unfamiliar tiles, Figure 3F, blue) elicited strong responses, followed by less familiar tiles (higher nslc, Figure 3F, yellow). Familiar tiles (nslc=1, i.e., tiles that had just been seen in the preceding trial) elicited almost no response (Figure 3F, red). The strong correlation between the neural responses, novelty, and familiarity can also be readily appreciated in individual trials (Figure 3G).

The reaction time was also a significant predictor for the neural responses recorded from this electrode (Figure 3E). However, the differences in neural responses signaling novelty and distinct degrees of familiarity *cannot* be explained by differences in reaction time. The differences in neural responses associated with novelty and familiarity persisted after reaction time equalization (see vertical dashed lines indicating equalized RT in Figure 3F).

The nslc predictor was statistically significant (p<0.01, GLM) in 45 gray matter electrodes (6.7% of the total, **Table S4A**, Figure 3H) and 32 white matter electrodes (6.5%, **Table S4B**, **Figure S5D**). **Figure S5** shows an example electrode located in the white matter whose responses correlated with novelty and familiarity. The majority of electrodes (82.2%) showed a positive correlation between the neural responses and nslc as illustrated in Figure 3E-G. The remaining electrodes (17.8%) showed a negative correlation, i.e., stronger neural responses for more familiar items. **Figure S6** depicts an example electrode located in the right pars opercularis showing a negative correlation between familiarity and neural responses.

Both the electrode in Figure 3E-H and the one in **Figure S6** revealed first-click as a significant predictor in addition to nslc, meaning that their responses not only reflected the familiarity gradient but also represented novelty. The electrodes that showed both first-click and n-since-last-click as significant predictors (20 electrodes) are denoted by red circles in Figure 3D and Figure 3H. Among these 20 electrodes, the signs of the t-statistic for the first-click and n- since-last-click predictors were consistent for 18 electrodes (16 positive and 2 negative). Only two electrodes exhibited opposite signs. These results indicate that novelty largely resembles extremely low familiarity in terms of the underlying neural responses.

### Neural signals show anticipation of the trial’s outcome

After seeing the 1^st^ tile, participants attempt to find the tile’s pair. If the 1^st^ tile’s pair was never encountered before, this is a random choice among the unseen tiles. If the tile’s pair is unfamiliar, recalling a match is error-prone and often leads to mismatches (Figure 2E). For highly familiar cases, participants can retrieve the correct location to find the match tile. Therefore, we asked whether the neural responses after exposure to the first tile and before seeing the 2^nd^ tile could predict successful retrieval.

Figure 4 shows the neural activity of an electrode located in the right lateral orbitofrontal cortex (arrow in Figure 4E), whose responses were predictive of successful retrieval. The GLM analysis indicated that match was a significant predictor of the neural responses (p<0.001, Figure 4A). This electrode showed stronger responses during match trials (Figure 4B, green) than during mismatch trials (Figure 4B, black). These differences can even be appreciated in single trials (compare Figure 4C versus Figure 4D). Of note, these differences are evident shortly after visualization of the first tile, with a peak at 500 ms after clicking the first tile, well before clicking the second tile, when the participant did not know for certain yet whether the trial would be a match or not. Thus, the strong neural differences between match and mismatch trials reflect the participant’s internal retrieval of the correct pairs’ locations.

**Figure 4.**
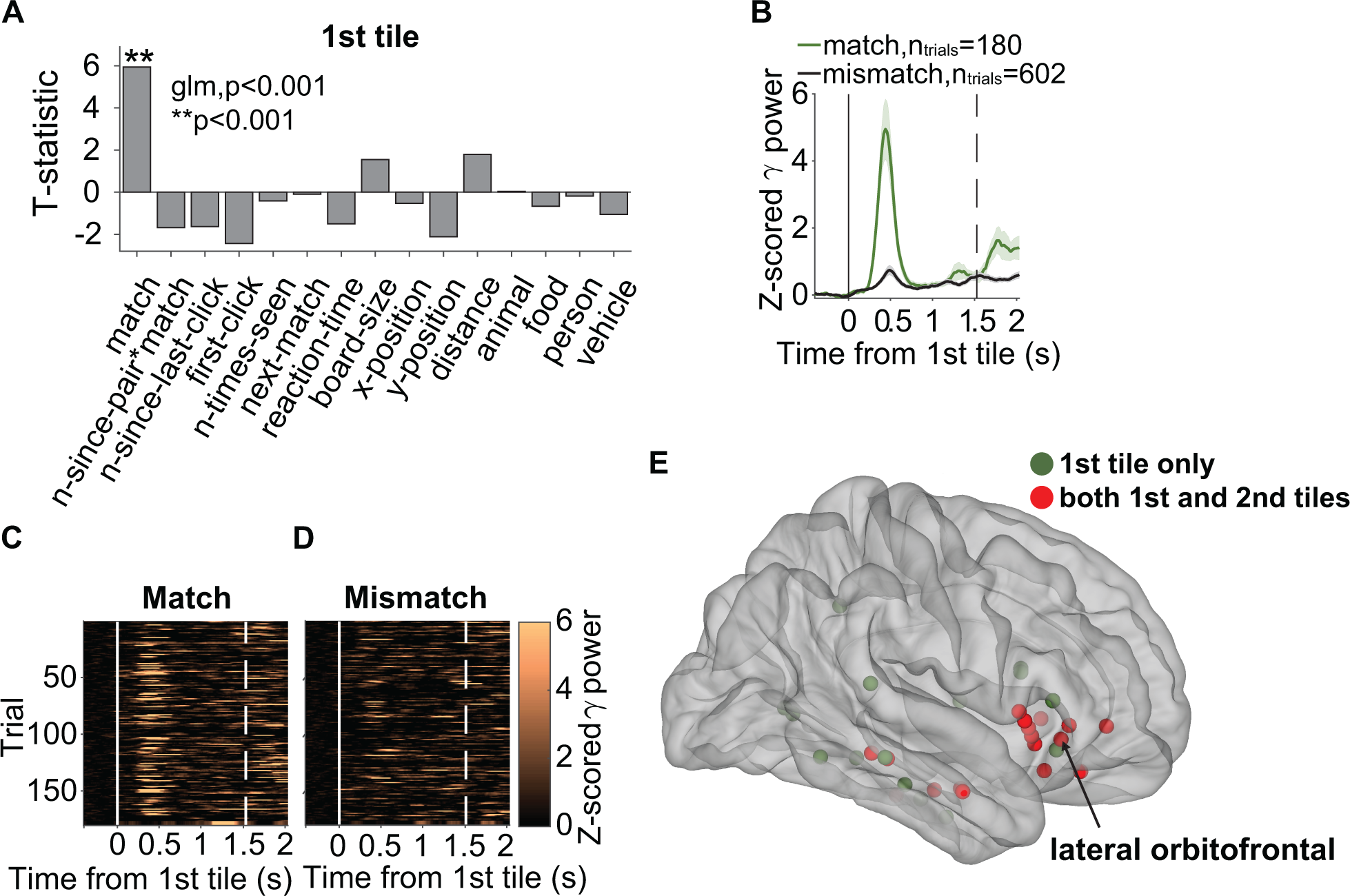
Neural signals predict correct retrieval Panels show an example electrode in the right lateral orbitofrontal cortex (see arrow in part **E**) and population locations in **E**. The format follows **Figure 3**. **A.** T-statistic of each predictor in the GLM analysis. Asterisks indicate statistically significant predictors for the neural signals. **B**. Z- scored gamma band power aligned to the 1^st^ tile onset (solid vertical line) for match trials (green) and mismatch trials (black). The vertical dashed line indicates the mean reaction time. Shaded error bars indicate s.e.m. **C**-**D**. Raster plots showing the gamma power in individual trials for match (left) and mismatch (right) trials. **E**. Locations of all electrodes where match was a significant predictor during the 1st tile only (green) and during both tiles (red). All electrodes were reflected on one hemisphere for display purposes.

Whether a trial was a match or not was a significant predictor of the neural responses for 32 electrodes in the gray matter (4.7% of the total, **Table S5A**, Figure 4E) and 30 electrodes in the white matter (6% of the total, **Table S5B**, **Figure S7**). For an example electrode in the white matter see **Figure S7**. In most cases (91%), neural activity was higher during match trials than during mismatch trials, as illustrated in Figure 4. The locations of all these electrodes, shown in Figure 4E (gray matter) and **Figure S7** (white matter), reveal that the majority were located in the lateral orbitofrontal (LOF) cortex, the medial temporal lobe, and the insula. The LOF cortex contained significantly more electrodes than expected by chance (p<0.01, **Methods**).

The peak in neural activity occurred at approximately 500 ms after the 1^st^ click (Figure 4B). As discussed in the previous section, the match predictor correlated with several other predictors (**Figure S4**). However, the GLM analysis shows that the match’s presence, but not other predictors, accounts for the neural responses (Figure 4A). To further establish this point, **Figure S8A** shows the responses of this same electrode, in the same format as Figure 4B, after equalizing the n-since-last-click (nslc) distributions for match and mismatch trials by subsampling the data. The same conclusions hold in this case. Furthermore, **Figure S8B** shows each match trial’s gamma power AUC versus the value of n-since-last-click and **Figure S8C** displays the same data from mismatch trials. The variable nslc did *not* account for the neural responses in either case (p>0.18, linear regression). Similar conclusions hold for the other predictors.

Figure S9 shows another example electrode located in the left middle temporal gyrus where the match was a significant predictor for the gamma band activity between the 1^st^ and 2^nd^ tiles. Similar to the LOF electrode in Figure 4, the gamma power during match trials was higher than during mismatch trials. However, the pattern of modulation in this electrode was sustained rather than transient (compare **Figure S9** versus Figure 4B-D). The change in gamma power was also evident in individual trials (**Figure S9B**-**C**). These observations suggest that the middle temporal and lateral orbitofrontal regions might be functionally distinct during memory retrieval.

In sum, these results indicate that even before the actual realization of whether a trial was a match or mismatch (i.e., before the onset of the 2^nd^ tile), there were distinct neural responses that were predictive of the trial’s outcome.

### Neural signals reflect the strength of memory retrieval

In addition to reflecting the outcome of a given trial (match versus mismatch), we considered the *n-since-pair* (nsp) predictor as a proxy for the degree of confidence or strength of memory retrieval. The smaller the nsp, the more recently *the tile’s pair* had been seen (Figure 2E-F). This predictor is different from n-since-last-click, which indicates how recently *the same tile*, rather than its pair, had been seen (Figure 1). We considered only match trials for this predictor (n-since-pair*match) because there was no successful retrieval of the tile’s pair in mismatch trials.

Figure 5 shows an example electrode in the left middle temporal gyrus (arrow in Figure 5E). In contrast with the electrode in Figure 4, both the match predictor and the nsp predictor were significant in the GLM analysis (Figure 5A). The t-statistic for nsp was negative, indicating a *decrease* in gamma band power for matching pairs that were more distant in memory. Indeed, responses were strongest for those tiles whose pairs had been seen less than 2 clicks ago (Figure 5B, red) and weakest when matching pairs had been seen more than 10 clicks ago (Figure 5B, purple). There was a negative correlation between the area under the curve (AUC) of the gamma band power and nsp (Figure 5C, p<0.001, linear regression). This correlation disappeared when considering mismatch trials (Figure 5D, p=0.66, linear regression), suggesting that the relationship between the neural signals and memory strength was contingent on successful retrieval.

**Figure 5.**
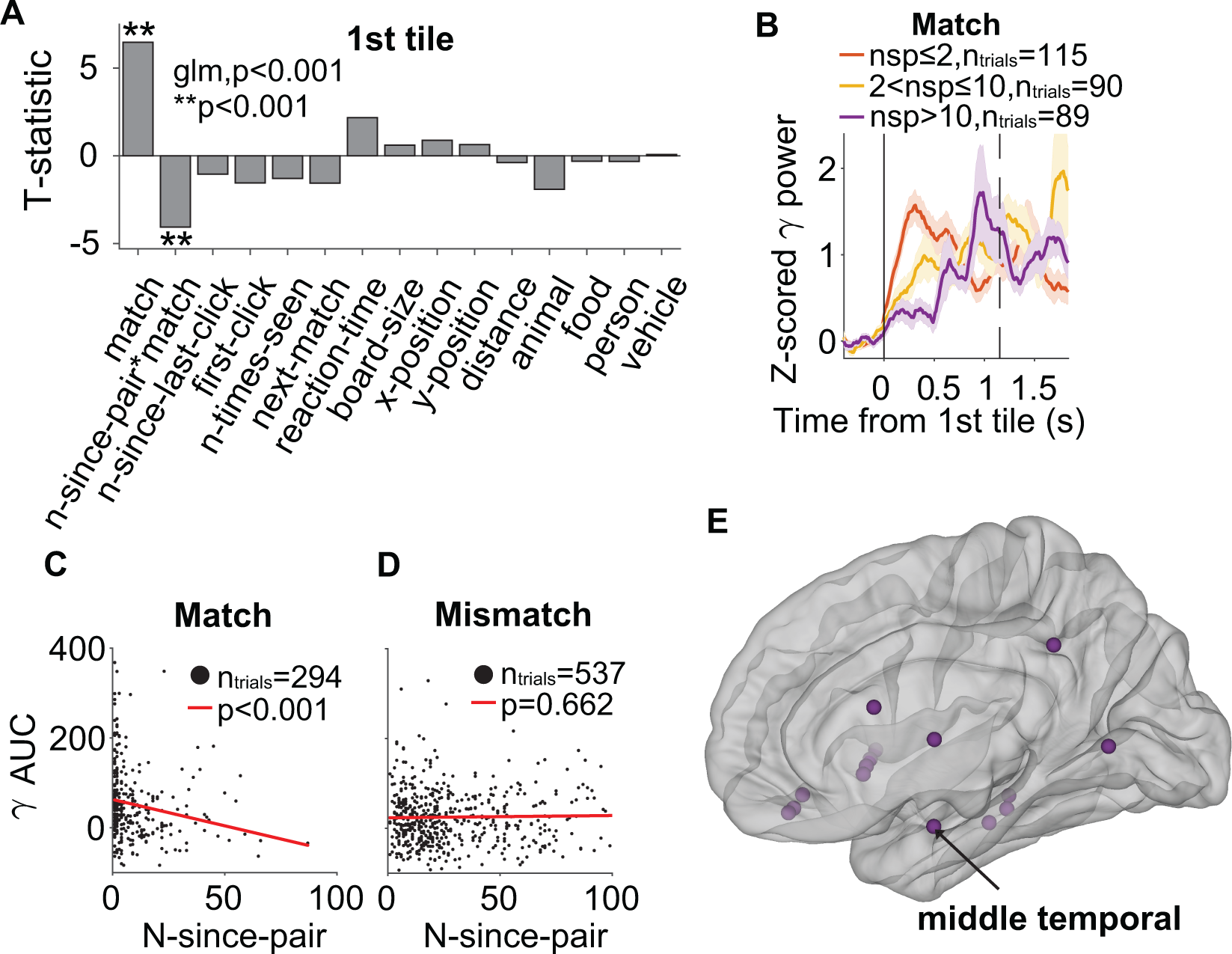
Neural signals reflect the strength of memory retrieval Panels show an example electrode in the left middle temporal gyrus (see arrow in part **E**) and population locations in **E**. **A**. T-statistic of each predictor in the GLM analyses. Asterisks indicate significant predictors for the neural signals. **B**. Z-scored gamma band power aligned to the 1^st^ file onset (solid vertical line) for match trials with small n-since-pair (nsp) (red, stronger memories), intermediate nsp (yellow), and large nsp (purple, weaker memories). The vertical dashed line indicates the mean reaction time. Shaded error bars indicate s.e.m. **C-D**. Scatter plots of the area under the curve (AUC) of the gamma band power as a function of nsp for match trials (**C**) and mismatch trials (**D**). Each dot represents one trial. Red lines show linear fits to the data. **E**. Locations of all electrodes where nsp was a significant predictor. All electrodes were reflected on one hemisphere for display purpose.

The nsp predictor was statistically significant (p<0.01, GLM) in 15 electrodes in the gray matter (2.2% of the total, **Table S6A**, Figure 5E) and 9 electrodes in the white matter (1.8% of the total, **Table S6B**, **Figure S10E**). For an example electrode located in the white matter, see **Figure S10**. Most of these electrodes showed a negative t-statistic, as in the example in Figure 5, and three electrodes (20%) showed the reverse effect (i.e., an increase in the neural signal for more distant associative memories). For most of these electrodes (73.3%), *match* was also a significant predictor, as illustrated by the example in Figure 5A, indicating that the neural signals encoded *both* successful retrieval *and* memory strength.

### Neural signals reflect feedback after the second tile

We have thus far focused on describing the responses elicited by the first tile in each trial. Next, we evaluated the neural responses triggered by the click of the second tile. We first asked whether novelty and familiarity were also encoded in the neural responses after the 2^nd^ tile. We built a separate GLM using the same 15 predictors except for n-since-pair*match (**Table 1**) to describe the AUC of the gamma power during one second after clicking the 2^nd^ tile. We excluded n-since-pair*match here because it would always be 1 during match trials (by definition, the first click was the pair of the second click). For the second tile, first-click was a significant predictor in 24 electrodes in the gray matter (3.6% of the total, **Table S7A**, **Figure S11E**) and 12 in the white matter (2.4% of the total, **Table S7B**). Thus, less than half the number of electrodes reflected novelty during the 2^nd^ tile compared to the first tile (cf. **Table S7A** versus **Table S3A** and **Table S7B** versus **Table S3B**). For the second tile, *n-since-last-click* was a significant predictor in 9 electrodes in the gray matter (1.3% of the total, **Table S8A**, **Figure S11D**) and 24 in the white matter (4.9% of the total, **Table S8B**), again, less than half the number of electrodes reflecting familiarity during the first tile (cf. **Table S8A** versus **Table S4A** and **Table S8B** versus **Table S4B**).

An example electrode in the LOF region whose responses correlated with familiarity after the 2^nd^ tile is shown in **Figure S11**. The LOF cortex contained significantly more electrodes than expected by chance (p<0.01, **Methods**). Among all the 9 electrodes where n-since-last-click was a significant predictor during the 2^nd^ tile, 7 electrodes also had first-click as a significant predictor (**Figure S11D-E**, red circles). In sum, novelty and familiarity of a tile were still encoded in the neural responses to the second tile, but to a lesser degree than during the responses to the first tile. This reduction may be due to the fact that two images were presented simultaneously, and the neural signals might reflect a weighted combination of the responses to each^64^. Moreover, for match trials, the information about the 2^nd^ tile does not need to be encoded in memory anymore to thrive in the task.

Among the 83 electrodes that had first-click as a significant predictor during the 1^st^ tile and the 36 electrodes during the 2^nd^ tile (including both gray and white matter), 15 electrodes (11 gray matter + 4 white matter) overlapped, i.e., first-click was a significant predictor during both the 1^st^ and the 2^nd^ tiles. Among the 77 electrodes that had n-since-last-click as a significant predictor during the 1^st^ tile and the 33 electrodes during the 2^nd^ tile (including both gray and white matter), 20 electrodes (5 gray matter + 15 white matter) overlapped, i.e., nslc was a significant predictor during both the 1^st^ and the 2^nd^ tiles. These electrodes may reflect general rather than specific novelty or familiarity mechanisms, irrespective of tile order, image content, or location.

Next, we asked whether the differences between match and mismatch trials were also manifested *after* the 2^nd^ tile was revealed, i.e., after the participant became explicitly aware of whether the trial was a match or not. Figure 6 shows an example electrode located in the left insula (see arrow in Figure 6E) where match was a significant predictor for the neural responses after the 2^nd^ tile (p<0.001, GLM, Figure 6A). The neural signals during match trials were larger than during mismatch trials (Figure 6B) and could be readily observed even in single trials (Figure 6C vs. **7D**).

**Figure 6.**
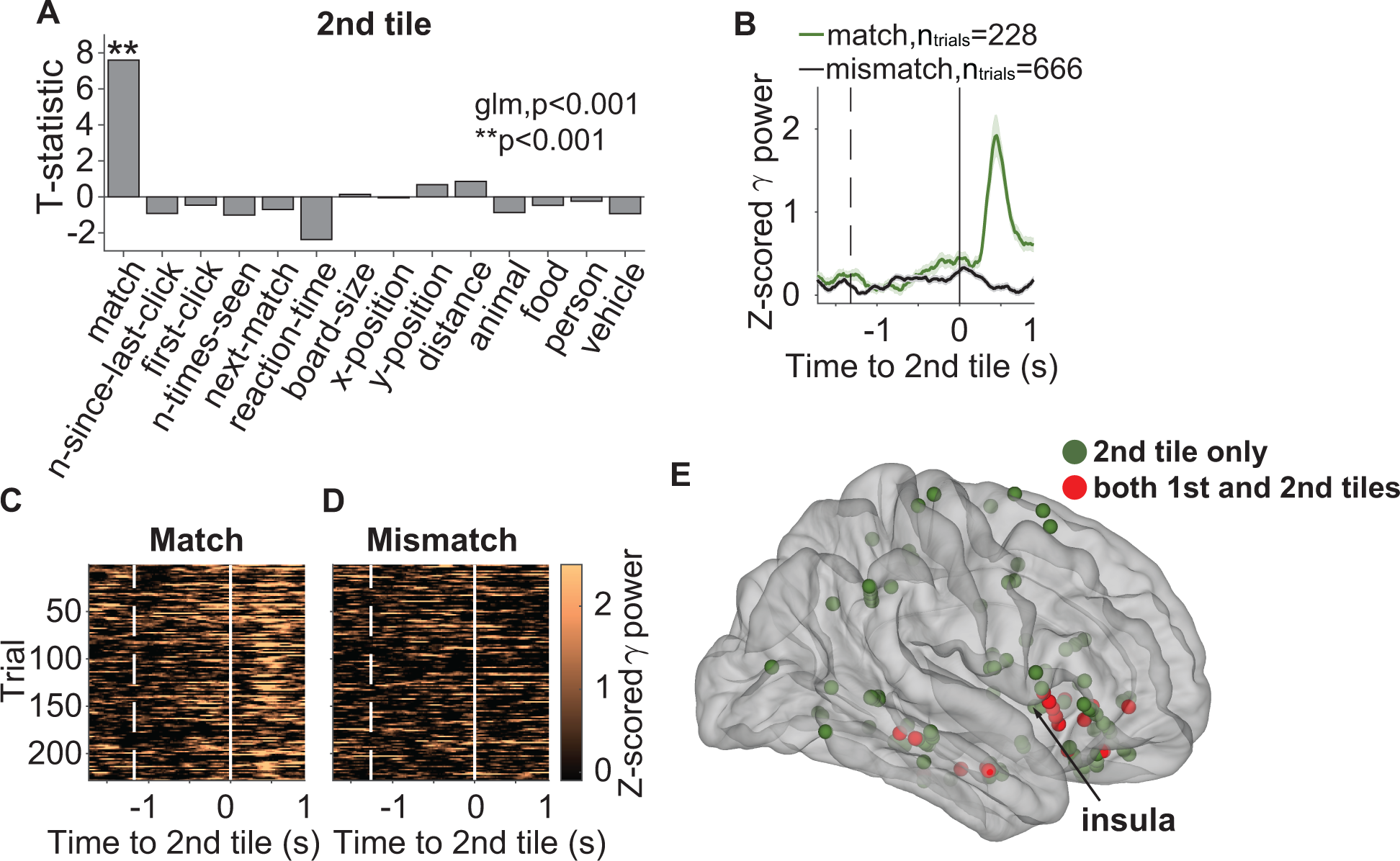
Neural signals after the second tile reflect correct retrieval Panels show an example electrode located in the insula (see arrow in part **E**) and population locations. **A**. T-statistic of each predictor in the GLM analyses for the responses after the 2^nd^ tile. Asterisks indicate significant predictors of neural signals. **B**. Z-scored gamma band power aligned to the onset of the 2^nd^ tile (solid vertical line) for match (green) and mismatch (black) trials. The dashed line indicates the mean onset of the 1^st^ tile. **C**-**D**. Raster plots showing the gamma band power in individual trials for match (left) and mismatch (right) trials. **E**. Locations of all electrodes where match was a significant predictor of neural responses after the 2^nd^ tile only (green) or during both tiles (red). All electrodes were reflected on one hemisphere for display purpose.

After the 2^nd^ tile, the match predictor was statistically significant (p<0.01, GLM) for 112 electrodes in the gray matter (16.6% of the total, **Table S9A**, Figure 6E) and 66 electrodes in the white matter (13.4% of the total, **Table S9B**, **Figure S12E**). For an example electrode in the white matter, see **Figure S12**. The locations of all these electrodes, shown in Figure 6E (gray matter) and **Figure S12E** (white matter), reveal that the majority were circumscribed to the LOF and the insula. The proportions of significant electrodes in both regions were higher than expected by chance (p<0.01, **Methods**).

There were 17 electrodes in the gray matter and 15 in the white matter where the match predictor was significant for both the 1^st^ and the 2^nd^ tiles. These electrodes represented 53.1% (gray matter) and 50% (white matter) of the electrodes that were significant according to the 1^st^ tile, and 15.2% (gray matter) and 22.8% (white matter) of the electrodes that were significant according to the 2^nd^ tile. These electrodes were located in the lateral orbitofrontal cortex, the medial temporal lobe, and the insula (Figure 4E, **S6E**, **7E**, and **S11E**, red circles). The electrode in **Figure S12** exemplifies such responses (compare the difference between match and mismatch before the onset of the 2^nd^ tile in **Figure S12B** versus Figure 6A). The electrode in **Figure S9** reveals a continuous enhancement for match trials after the 1^st^ tile that was sustained and continued after the onset of the 2^nd^ tile.

### A machine learning classifier could predict matches in single trials

We evaluated whether the neural responses within a brain region could predict if a trial was a match or a mismatch (**Methods**). For this analysis, we considered only those brain regions with more than 12 electrodes combining all participants (**Methods**). Figure 7 shows the average decoding performance of an SVM classifier after 200 iterations of 5-fold cross-validation. At each iteration, the SVM consisted of a binary classifier (match versus mismatch). The predictors were the PCA features extracted from the concatenated neural responses of electrodes within a particular brain region. The superior parietal gyrus and insula exhibited decoding accuracy above chance (p<0.01, **Methods**) during the 1^st^ tile (Figure 7A). The lateral orbitofrontal and middle temporal cortex also showed accuracy above chance (>60%, Figure 7A), albeit not statistically significant. As expected based on the responses of individual electrodes, the decoding accuracy was higher after the 2^nd^ tile compared to the 1^st^ tile. After the 2^nd^ tile, multiple brain regions showed accuracy above chance (p<0.01, Figure 7B). The lateral orbitofrontal cortex, insula, middle temporal gyrus, and pars opercularis showed the highest accuracy (>75%). Similar results were obtained when subsampling 12 electrodes during each iteration (**Figure S13**).

**Figure 7.**
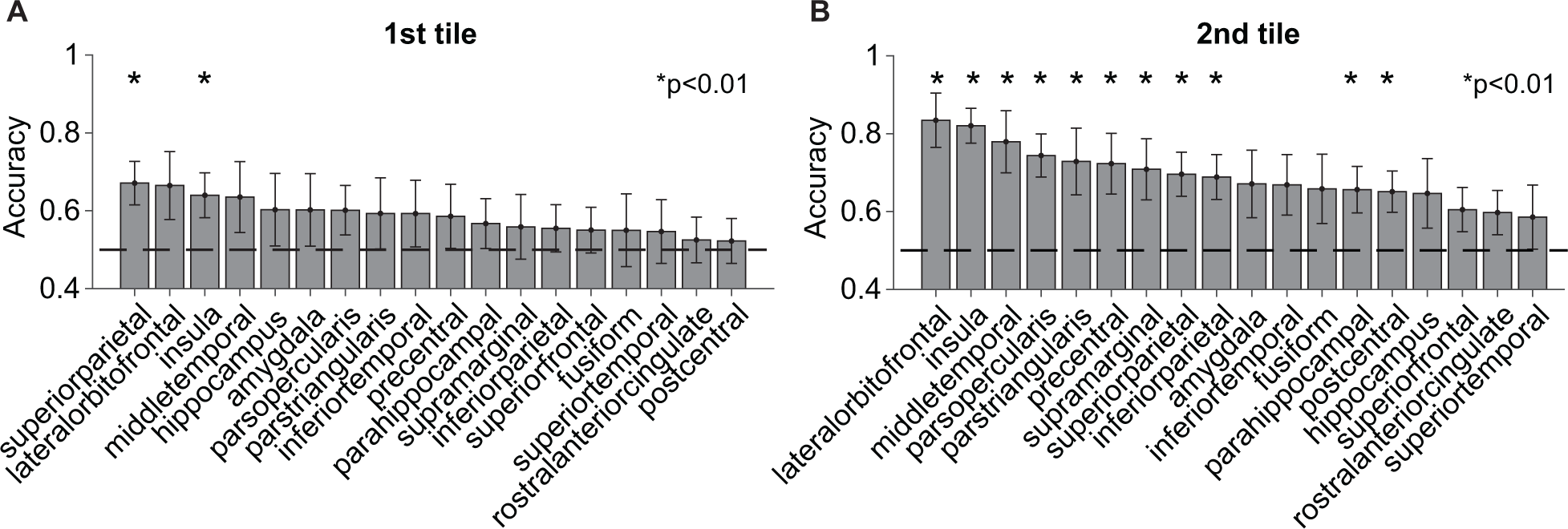
Machine learning decoding of match versus mismatch for all channels within each brain region Average decoding accuracy for each brain region (**Methods**) using neural responses after the 1^st^ tile (**A**) or after the 2^nd^ tile (**B)**. Brain regions are ordered from higher to lower average decoding accuracy. The dashed horizontal line indicates chance accuracy. Asterisks denote significant decoding accuracy above chance (⍺=0.01). All error bars indicate SD (n=1,000 iterations).

### A computational model provides a first-order approximation to the behavioral and neural measurements

To further understand the mechanisms at play during the task, we built a computational model that focused on the storage and retrieval of information (Figure 8, **Methods**). The computational model consists of a Hebbian attractor neural network with all-to-all connectivity.

**Figure 8:**
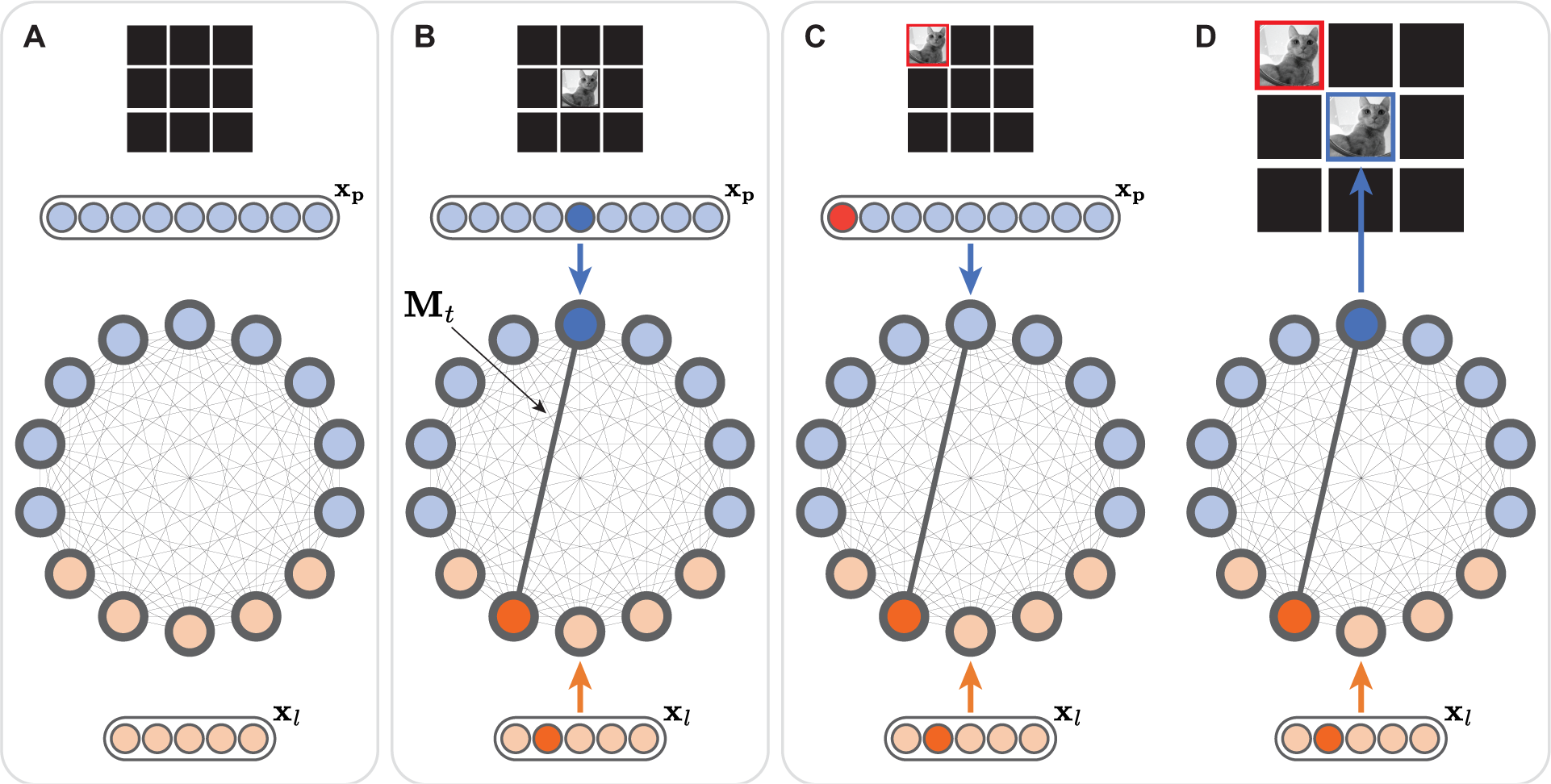
Hebbian attractor model architecture and operating regimes **A.** Schematic representation of the model architecture used for the 3x3 grid. The 9 blue units encode position (***x***_*l*_), while the 5 orange units represent the image label (***x***_*l*_). The black lines between units illustrate the Hebbian weights *M*_*t*_ in the attractor network. **B.** Learning regime. In this example, the model represents a cat (label=2) at position=5. **C, D.** Inference regime. In this example, the model is tasked with matching the cat (label=2) observed at position=1. Only the label information is provided to the model in the inference regime. The model’s updates (**Methods**) lead to the unit representing position=5 to exhibit the highest activity (**D**), thereby determining the corresponding tile to be clicked. The darker color indicates stronger activation of the corresponding units. The red color indicates the tile to match (which is unavailable for clicking, **Methods**) and its corresponding positional unit.

The units are divided into position units (the number equaling the number of tiles on the board) and label units (the number equaling the number of images on the board) (Figure 8A). The model has two main modes of operation: learning (Figure 8B), and inference (Figure 8C-D). After the first click, the model receives as input the label of the tile and its position. The activity of each unit evolves over time based on the external input and the weighted input from other units followed by a rectifying non-linearity and normalization (**Methods**, Equation 1). Concomitantly, the weights are updated in a Hebbian manner (**Methods**, Equation 2). During inference, the model selects the position unit with the maximum activation for the second click. The model proceeds in this manner until all matches have been found.

We computed the same performance evaluators from Figure 2A,B,D,F for the model. We did not compute the metrics for Figure 2C,E because the model chooses the first tile randomly among the available tiles (**Methods**). We defined the reaction time as the number of steps needed for the selected unit to reach 0.9 of its maximum value (**Methods**). The number of clicks per tile increased with the board size, approximating the participants’ behavior (compare Figure 9A versus Figure 2A). The reaction time for the model was longer for mismatch trials than for match trials for all board sizes (p<0.001, compare Figure 9B versus Figure 2B). The nslc value increased with board size and was much larger for mismatch trials compared to match trials, consistent with the participants’ behavior (p<0.001, compare Figure 9C versus Figure 2D). Similarly, the nsp value increased with board size and was also significantly larger in mismatch trials compared to match trials for all board sizes (p<0.001, compare Figure 9D versus Figure 2F). To investigate the model’s inner workings, we defined two metrics based on the unit activations. First, to compare with the match related signals in Figure 3, we computed an overall maximum energy (**Methods**, Equation 3). This maximum energy was smaller for trials with nslc=1 (p<0.001, Figure 9E), reflecting a strong correlate of memory for recently seen tiles (compare to Figure 3B and especially Figure 3F). Second, we defined a confidence metric by assessing the relative activation for the strongest unit with respect to the other units during the inference step (**Methods**). The confidence metric was significantly larger for match trials compared to non-match trials (p<0.01, Figure 9F), which was qualitatively similar to the neural responses described in Figure 5.

**Figure 9:**
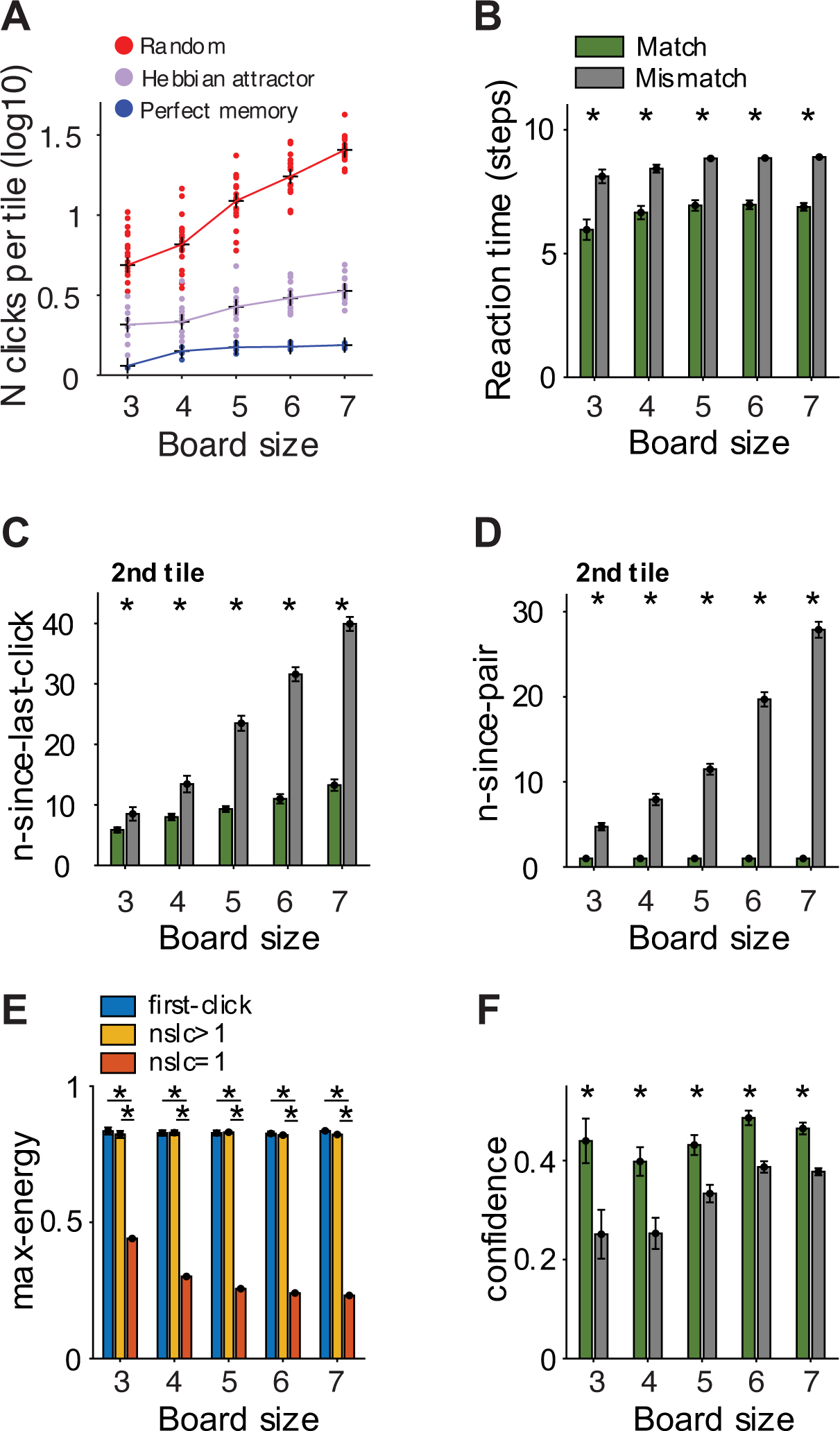
Hebbian attractor model captures behavior and neural measures **A**. Number of clicks per tile (log scale) as a function of board size for random simulation model (red, n=20), perfect memory simulation model (blue, n=20), and Hebbian attractor (purple, n=20) (**Methods,** compare to **Figure 2A**). **B**. Reaction times of the model (Methods) for match (green) and mismatch (gray) trials for different board sizes (compare to **Figure 2B**). **C**, **D**. Average n-since- last-click (**C**) and n-since-pair (**D**) for the 2nd click for each board size (compare to **Figure 2D**, **F**. **E**). Max-energy for novel tiles (blue), unfamiliar tiles (n-since-last-click>1, yellow) and familiar tiles (n-since-last-click=1, red, compare to **Figure 3**). **F**. Model confidence for match (green) and mismatch (gray) trials for different board sizes (**Methods**). The model confidence in match trials was larger than in mismatch trials (compare to **Figure 5**). All asterisks denote significant difference between match and mismatch trials (permutation test, 5,000 iterations, α=0.01). All error bars indicate s.e.m. (n=20).

## Discussion

In this study, we investigated the dynamics of neural signals during a natural memory task where participants engaged in a classical card-matching game. Participants performed the task well, slightly worse than expected by a perfect memory model (Figure 2A), showing increased reaction times during mismatch trials (Figure 2B) as well as decay of memory traces with time since encoding (Figure 2C-F).

Many studies have focused on studying memory for a list of sequentially presented stimuli ^1,7,12,19,24,65,66^ or examined memory in naturalistic or real-world scenarios at the behavioral level ^67–71^. The task introduced here strikes a balance between these two approaches, presenting a more realistic and complex setting that involves associative and non-associative memories within the same task and introducing task-dependent complexity compared to word lists. Yet, our task allows for a high level of control over stimulus timing and experimental parameters that are difficult to achieve when studying memory in real-world scenarios.

Complex and natural tasks necessarily depend on the interplay of multiple intercorrelated variables. To tame the complexity of these different variables, we used generalized linear models (GLMs) to quantitatively assess the influence of distinct predictors on the neural responses. Through these GLM analyses, we could characterize neurophysiological responses that were largely governed by individual predictors after accounting for the correlations among predictors (**Figure S4**). While focusing on any one predictor, these results were corroborated by subsampling the data to equalize other predictor variables that could potentially affect the neural responses. The extensive sampling of brain regions, including neural activity from more than 1,000 electrodes across 20 participants (**Figure S2-S3**), allowed us to track neural responses during each of the steps required in the task with broad brain coverage.

The first step to encode tile information is to correctly determine whether a tile is novel or not. We refer to the ability to detect novelty and familiarity as non-associative recognition memory^16,24^. Assessment of novelty and familiarity orchestrate strategies of encoding and maintenance of information in memory^1,72^. We found strong neural responses that signal novelty and familiarity (Figure 3, **Figure S5-6**, **S10-11**, **Table S3**, **S4**, **S7** and **S8**). Several electrodes responded both to novelty and familiarity, showing similar responses to novel and highly unfamiliar items (Figure 3F-G and **Figure S5B-C**). The lateral orbitofrontal cortex, the pars opercularis, and the medial temporal lobe contained a preponderance of electrodes signaling novelty and familiarity. These responses are reminiscent of novelty and familiarity signals that have typically been described in tasks involving a sequence of images presented with occasional repetitions^6,16,18,66^ but are not restricted to the medial temporal lobe. While some electrodes may also encode information about the image content (Figure 3E), most of the electrodes signaling novelty and familiarity were not content-specific, consistent with the observation that neurons involved in memory formation are rarely sharply tuned to particular sensory features^28^.

After seeing the first tile in a trial, participants could estimate whether they remembered the location of its pair and, thus, whether they would get a match or not. Participants link the first tile with its pair to internally predict where to click next. Indeed, neural responses strongly reflected not only these predictions (Figure 4, **Figures S7**-**9**) but also the internal estimate of the memory strength or the confidence of these predictions (Figure 5). Even though participants could not know for sure yet whether the trial would be a match or not, the neural signals exhibited large differences between match and mismatch trials after the first tile and before the revelation of the second tile. These differences were either transient (Figure 4) or sustained (**Figure S9**). We speculate that transient increases in activity during match trials might signal the sudden realization and high confidence about the trial outcome (match or mismatch). In contrast, sustained responses may correspond to the active retrieval processes. It is possible that sustained activity could arise due to averaging across temporally shifted transient activities from individual neurons^73^; however, it has been reported that single neurons in the hippocampus can also show sustained firing rate increase for successful associative retrieval^74^. Extensive work has documented the importance of the hippocampus and surrounding structures in the MTL in associative memory (e.g.,^16,26,74,75^). In addition to the MTL, the current results show that other areas, such as the lateral orbitofrontal cortex, also play a critical role in associative recall.

Several studies have highlighted the potential for attractor-based models to characterize working memory processes^47,49,76,77^. Here we show as a proof-of-principle that a simple instantiation of an attractor-based neural network model can qualitatively capture the properties of human participants both at the behavioral level (Figure 9A-D) and also at the neural level (Figure 9E-F). This basic neural network architecture can be readily linked to a visual neural network backbone to further examine the underlying representation of visual signals in working memory. Additionally, the model could be extended to examine even more complex tasks that involve multiple-way associations and dynamic changes in the structure of the environment over time. The high temporal resolution, extensive spatial sampling, and computational models provide an opportunity to characterize the dynamics of complex naturalistic tasks. These observations provide initial steps to further our understanding of how different components of encoding and retrieval interact during the formation of natural memory events.

## Methods

### Task paradigm

Participants performed our implementation of the classical memory matching game (Figure 1**, Movie S1**). The game involves remembering the location and content of a set of tiles to find all the matching pairs. A square board containing *n* × *n* tiles was shown throughout each block. In the beginning, all tiles were shown in black. In each trial, participants chose one tile, and then a second tile, by clicking on them in a self-paced fashion. Upon clicking, the tile revealed a common object like a cat or an indoor scene like a kitchen. At the end of each trial, either the two tiles revealed the same content (match) or not (mismatch). If the tiles matched, then the two tiles turned green 1,000 ms after the second click, and the two tiles could not be clicked again for the remainder of the block. If the tiles did not match, they turned black 1,000 ms after the second click and could be clicked again in subsequent trials. When all tiles turned green, i.e., all matches were found, the block ended, and another block began. During each block, the map between positions and objects was fixed. The game always started with a block of size 3×3 and progressed to more difficult blocks (4×4, 5×5, 6×6, and finally 7×7). Blocks with an odd number of tiles (3×3, 5×5, and 7×7) contained one distractor object (a human face) with no corresponding pair. For each block except the 3×3 board, there was a limit for the total time elapsed (2 minutes for 4×4, 3.3 min for 5×5, 4.8 min for 6×6, and 8.2 min for 7x7). If a participant did not complete a block within the time limit, the block ended, and a new, easier block started by reducing the board size *n* by 1, except when *n*=7, where it was reduced by 2. Conversely, when participants successfully completed a block with a board of size *n* within the allotted time limit, they moved on to a more difficult block by increasing *n* by 1. When participants completed an *n*=7 block, they performed further *n*=7 blocks. There was no image repetition across blocks.

All the images were from the Microsoft COCO 2017 validation dataset ^78^ and were rendered in grayscale and square shape. We included a balanced number of pictures from 5 categories: person, animal, food, vehicle, and indoor scenes. All the images were rendered on a 13-inch Apple MacBook Pro laptop. The size of each tile was 0.75×0.75 inches (approximately 2x2 degrees of visual angle, dva) and the separation between two adjacent tiles was 0.125 inch (0.33 dva) for board size *n*=7 and 0.25 inch (0.67 dva) for the others. The game implementation was written and presented using the Psychtoolbox extension ^79,80^ in Matlab_2016b (Mathworks, Natick, MA).

### Epilepsy participants and recording procedures

We recorded intracranial field potentials from 20 patients with pharmacologically intractable epilepsy (12-52 years old, 9 female, **Table S1**) undergoing monitoring at Boston Children’s Hospital (Boston, US), Brigham and Women’s Hospital (Boston, US), and Xuanwu Hospital (Beijing, China). All recording sessions were seizure-free. All patients had normal or corrected-to-normal vision. The study protocol was approved by each hospital’s institutional review board. Experiments were run under patients’ or their legal guardians’ informed consent. One patient at Brigham and Women’s Hospital (BWH) was implanted with both stereo encephalography (sEEG) and electrocorticography (ECoG) electrodes, while all other patients had only sEEG electrodes (Ad-tech, USA; ALCIS, France). Intracranial field potentials were recorded with Natus (Pleasanton, CA) and Micromed (Italy). The sampling rate was 2048 Hz at Boston Children’s Hospital (BCH), 512 Hz or 1024 Hz at BWH, and 512 Hz at Xuanwu Hospital (XWH). Electrode trajectories were determined based on clinical purposes for precisely localizing suspected epileptogenic foci and surgically treating epilepsy^81^.

### Eye tracking procedures

Ten non-epilepsy healthy participants (23-35 years old, 9 female) performed the same task while their eye movements were tracked with the EyeLink 1000 plus system (SR Research, Canada) at a sampling rate of 500 Hz. The task paradigm was the same as the one for epilepsy participants except that, before each block began, participants fixated on a center cross to ensure that the EyeLink eye-tracking system was well-calibrated. Otherwise, a re-calibration session ensued. The task was presented on a 19-inch CRT monitor (Sony Multiscan G520), and participants sat about 21 inches away from the monitor screen. The tile size was 1x1 inches (approximately 2.7x2.7 degrees of visual angle) as appeared on the screen. The study protocol was approved by the institutional review board at Boston Children’s Hospital, and each participant completed the task with informed consent. All participants had normal or corrected-to-normal vision. All participants completed 16 blocks.

### Behavioral analyses

We created two computational models to simulate behavior assuming perfect memory or no memory (chance performance, Figure 2A). The perfect memory model remembered all revealed tiles without forgetting. The random model simulated random clicking. We calculated the reaction time (RT, time between two clicks in a trial), n-since-pair (number of clicks since the last time when a tile’s matching pair was seen), n-since-last-click (the number of clicks since the same tile was clicked), and n-times-seen (number of times the same image had been seen). For n-since-pair and n-since-last-click, we excluded trials in which any tile was seen for the first time, i.e., when a tile’s matching pair had never been revealed, or there was no previous click. We compared these variables for match and mismatch trials at each board size (Figure 2, permutation test, 5,000 iterations, α=0.01). We defined random matches as a match trial where the second tile had never been seen before; such trials were excluded from both the behavioral and neurophysiological analyses. We used the F-test for linear regression models to assess whether RT, n-since-pair, n-since-last-click, and n-times-seen significantly covary with board size. The linear regression models’ predictors were these four behavioral parameters and the dependent variable the board size. We created separate models for match and mismatch trials and 1^st^ and 2^nd^ tiles.

### Electrode localization

Electrodes were localized using the iELVis^82^ toolbox. We used Freesurfer^83^ to segment the preimplant magnetic resonance (MR) images, upon which post-implant CT was rigidly registered. Electrodes were marked in the CT aligned to preimplant MRI using Bioimage Suite^84^. Each electrode was assigned to an anatomical location using the Desikan-Killiany^85^ atlas for subdural grids or strips or FreeSurfer’s volumetric brain segmentation for depth electrodes. For white matter electrodes, we also reported their closest gray matter locations. Out of 1,750 electrodes in total, we included 676 bipolarly referenced electrodes in the gray matter (**Figure S2, Table S2**) and 492 bipolarly referenced electrodes in the white matter (**Figure S3, Table S2**). Five hundred and eighty-two electrodes were not considered for analyses due to bipolar referencing, locations in pathological sites, or electrodes containing large artifacts. Electrode locations were mapped onto the MNI305 average brain via affine transformation^86^ for display purposes (e.g., **Figure S2- S3**).

### Preprocessing of intracranial field potential data

Bipolar subtraction was applied to each pair of neighboring electrodes on each shank of depth electrodes or subdural grids/strips^87^. A zero-phase digital notch filter (Matlab function “filtfilt”) was applied to the bipolarly subtracted broadband signals to remove the line frequency at 60 Hz (BCH, BWH) or 50 Hz (Xuanwu) and their harmonics. For each electrode, trials whose amplitudes (Voltagemax-Voltagemin) were larger than 5 standard deviations from the mean amplitude across all trials were considered potential artifacts and discarded from further analyses ^88^. For the first tile, the time window for artifact rejection was from 400 ms before the click until 1 second after the average RT. For the second tile, the time window was [400 ms + average RT] before the second click until 1 second after the second click. Across all electrodes, we rejected 1.75% of all trials for the 1^st^ tile and 1.73% for the second tile.

### Time-frequency decomposition

The gamma band (30-150 Hz) power was computed using the Chronux toolbox^89^. We used a time-bandwidth product of 5 and 7 leading tapers, a moving window size of 200 ms, and a step size of 10 ms^90^. For each trial, the power was normalized by subtracting the mean gamma band power during the baseline (400 ms before 1^st^ tile) and dividing by the standard deviation of the gamma power during the baseline. For all the participants, there were more mismatch than match trials. In the raster plots, we subsampled the mismatch trials, keeping those trials whose reaction times were closest to the mean reaction time of match trials. All random matches were excluded from analyses.

### Generalized linear model

We used generalized linear models (GLM)^91,92^ to analyze the relationship between the gamma band power and behavioral parameters. We used two different GLMs, one using neural responses between the 1^st^ and the 2^nd^ tiles and the other using neural responses after the 2^nd^ tile. For the first GLM, the time window started when the 1^st^ tile was clicked and ended at a time corresponding to the 90th percentile of the distribution of reaction times (time difference between the 1^st^ and the 2^nd^ click, Figure 1A) for each participant. This criterion was a reasonable tradeoff between minimum overlap with responses after the 2^nd^ tile and the maximum amount of information captured. For the second GLM, we used 1 second after the 2^nd^ tile click as the analysis window. The response variable to be fit by the GLM analyses was defined as the area under the curve (AUC) of the gamma band power over the specified time windows. **Table 1** describes the behavioral parameters that were considered as predictors in the models.

We performed a multicollinearity analysis to assess the presence of highly correlated predictors that could impair the model’s performance^93,94^. We calculated the variance inflation factor (VIF) for each predictor to detect the presence of multicollinearities. A VIF of 1 indicates that there is no correlation with other predictors. The larger the VIF, the higher the correlation. A VIF greater than 5 indicates a very high correlation that could significantly harm the model’s performance. For all participants in our analysis, the VIFs of all predictors were smaller than 3 (**Figure S4C-D**).

For n-since-pair, we included the interaction term between this predictor and match (n- since-pair*match) to test the hypothesis that when a trial was a match, the strength of the neural response after the first tile was modulated by how recently the tile’s matching pair was seen for the last time. The neurophysiological responses confirmed that this is a reasonable way to model the data (Figure 5B). We represented image categories as predictor variables in the GLM by including four out of the five categories (animal, food, vehicle, and person). We dropped the “indoor” category to avoid falling into the “dummy variable trap”^95^. For each predictor, we calculated the parameter estimate (beta coefficient) from the least mean squares fit of the model to the data, the t-statistic (beta divided by its standard error), and the p-value to test the effect of each predictor on the neural responses. A beta coefficient or t-statistic of zero indicated that the predictor did not affect the neural responses. A predictor was considered statistically significant if the GLM model differed from a constant model (p<0.01) and the p-value for that predictor was smaller than 0.01.

To determine if any brain region contained significantly more electrodes than expected by chance considering any GLM predictor, we randomly sampled the same number (n) of electrodes as those where that predictor was significant, from the total electrode population (separately for gray and white matter electrodes) for 5,000 iterations. Taking the *match* predictor for gray matter electrodes as an example, from the total of 676 gray matter electrode locations, we randomly sampled 32 electrodes (the number of match-significant gray matter electrodes) 5,000 times, and calculated the p-value of any location, say insula, as the number of times when n for insula and match-significant was less than n for insula sampled. If p<0.01, that brain region was considered to have significantly more electrodes than expected by chance.

### Decoding of match

We used a machine learning decoding approach to evaluate whether the neural responses from a given brain region could predict whether the trial was a match or a mismatch. For this analysis, we selected only those brain regions in which we had at least 12 electrodes. We used two different decoders for each brain region, one using neural responses between the 1^st^ tile and 800 ms after the 1^st^ tile click, and the other using the neural responses between the 2^nd^ tile and 800 ms after the 2^nd^ tile click. We performed 200 iterations of 5-fold cross-validation for each brain region and tile to split the trials into independent train and test sets (1,000 splits in total). We concatenated the neural responses of electrodes within the same brain region, using two different approaches: (i) taking all electrodes in each brain region, and (ii) randomly subsampling 12 electrodes at each iteration. The number of match and mismatch trials was normalized by random subsampling. To reduce the dimensionality of the neural responses, we used principal component analysis (PCA). The PCA parameters were computed using only the training data, and we selected the number of components that could explain 70% of the training neural responses variance. We used support vector machines (SVM) with a linear kernel for the binary classification (match or mismatch), using as the model inputs the PCA features computed from the neural responses. We followed the same procedure to test whether the classification performance was above chance, but we randomized the response variable (match or mismatch) at every iteration. We calculated the p-value as the number of times when the accuracy using random labels was above the average accuracy using the actual labels. If p<0.01, neural responses of that brain region could predict match and mismatch above chance.

### Computational model

We developed an attractor network model consisting of a fully connected recurrent network with the number of units *n* equal to the number of tiles in the grid plus the number of different images. For example, the model for the 3x3 board shown in **Fig 1A** was an attractor network with *n*=3x3+5=14 units (Figure 8).

The units in the network were designed to model “where” and “what”, i.e., position and image labels. Let ***x***_*p*_be a vector of length equal to the number of tiles in the grid, ***x***_*l*_ be a vector of length equal to the number of different images in the grid, and ***x*** denote the concatenation [***x***_*p*_, ***x***_*l*_] (Figure 8A). The input to the network is ***x***. Each entry in ***x***_*p*_ and ***x***_*l*_ can take the values - 1, 0, or 1. The state of the network at time *t* is denoted by the vector ***h***_*t*_ = [***p***_*t*_, ***l***_*t*_] of size *n*, where ***p***_*t*_ and ***l***_*t*_ are the vectors of activations of the position and label units, respectively. Each entry in ***h***_*t*_ is a scalar value. The units in the network are connected in an all-to-all fashion and the matrix ***M***_*t*_ indicates the weights at time *t* (***M***_#_ ∈ ℝ^*n*×*n*^).

The network stores memories in both persistent activities (active representations) and weights (silent representations)^49^. In contrast to the approach in ref.^49^, which incorporates a bottleneck in the model to restrict its capacity, our model is devoid of any such bottleneck. Given an input ***x*** at time *t*, the network state and weights were updated similarly to ref.^96^, according to:

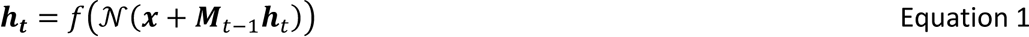

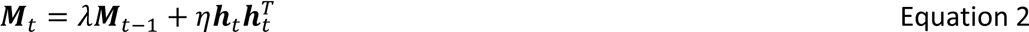

Here *f*(⋅) is the LeakyReLU activation function and 𝒩(⋅) is activation normalization. λ and η represent a decay rate for the previously stored memories and the learning rate for new memories, respectively. In line with ref.^96^, activation normalization is expected to make the network more robust to the choice of the decay and learning rates. We note that the Hebbian learning is computed on the state of the network ***h***_*t*_ rather than on the input ***x***. This means that the update of the memory matrix ***M***_*t*_ is influenced by the interference between active and silent representations, thus limiting the network capacity. The hyperparameters were chosen by fitting the number of clicks per tile of the model to the participants’ number of clicks per tile (Figure 2A). The results presented in this paper were obtained with λ = 0.6 and η = 0.9. Before the start of each board, the network weights were initialized uniformly at random in [0,1], while the state of the network was initialized to 0. Changes in weights in neural networks are often interpreted as structural modifications to synaptic strengths. However, given the time scales involved in working memory tasks such as the one studied here, changes in ***M***_*t*_ are more likely to reflect transient biophysical mechanisms such as synaptic facilitation rather than permanent structural synaptic modifications.

The model operates in two distinct regimes, which we refer to as *learning* (Figure 8B) and *inference* (Figure 8C-D). For each trial, the 1^st^ tile was chosen at random among the available tiles. To simulate the task, for each trial the model performs learning→inference→learning. First, the model represents the position and label of the 1^st^ tile. Second, the model performs inference on the label of the 1^st^ tile. At the end of the inference regime, the most active neuron in ***p***_*t*_ determines which tile to click (Figure 8D). Last, the model learns the position and label of the 2^nd^ tile.

During learning (Figure 8B), the corresponding position entry of ***x***_*p*_ is set to 1 and all other units are set to -1. Similarly, the corresponding label entry of ***x****l* is set to 1 and all other units are set to -1. The network dynamics goes through 10 steps according to the two equations above. During inference (Figure 8C-D), the corresponding label of ***x***_*l*_ is set to 1 and all the other units are set to 0. All the units of ***x***_*p*_ corresponding to the available tiles are set to 0, while the ones corresponding to the unavailable tiles (those that have already been matched or already clicked in that trial) are set to -1. The network dynamics goes through 10 steps according to Equations 1-2. After these 10 steps, we select the unit with the maximum activation within the units of ***x***_*p*_ corresponding to available tiles. If the second tile is a match, then those two tiles become unavailable in the next trials. The weight matrix ***M***_*t*_, however, continues to include all the connections among all the units. The model proceeds until all tiles have been matched.

The number of clicks per tile, n-since-last-click and n-since-pair click for the 2^nd^ tile were calculated for the model and compared to the participants’ behavior (Figure 9A-D). To compute a proxy for the reaction time in the model, we used the same approach as in ref.^49^, whereby the unit in ***x***_*p*_ with the strongest activation during the inference time was selected and the reaction time was computed as the number of steps the unit takes to reach 0.9 of its maximum value.

To compare the inner workings of the model with the neural data, we defined two new metrics based on the unit activations. First, we defined the *max-energy* metric computed during the 1^st^ learning phase of each trial, in analogy to the memory signals in Figure 3. The energy of the network was computed as:

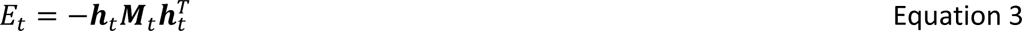

Min-max normalization was applied to the energy in each trial, and the maximum value in each trial was reported. The max-energy metric is shown in Figure 9E. Second, we defined a *confidence* metric that reflected the evidence for a match in a given trial, in analogy with the predictive signals shown in Figure 5. The confidence metric was defined by selecting the strongest activation in ***p***_*t*_during inference, subtracting the mean value of ***p***_*t*_, applying min-max normalization to the difference, and then taking the maximum over time *t* in each trial. The confidence metric is shown in Figure 9F.

### Data availability

We share all the deidentified psychophysics data, electrode location information, and neural data, together with all the code generated to model and analyze the data through the following public site: https://klab.tch.harvard.edu/resources/HowToGetAMatch.html

## Author contributions

The task was designed by YX and GK. All the data were collected by RJW (XWH) and YX (BWH and BCH) with the help of PHW (XWH) and DW (BWH). GGZ (XWH), YZS (XWH), CRG (BWH), JRM (BCH), and SS (BCH) performed the surgeries on the patients. All the data were curated by YX and analyzed by YX and PSL, with frequent discussions with GK. The computational model was developed by RS, with frequent discussions with GK. The manuscript was written by YX, PSL, RS, and GK and approved by all authors.

## Acknowledgments

This work was supported by the McKnight Foundation, NIH Grant R01026025, and NSF Grant CCF-1231216.

## Supplementary Materials

**Figure S1.**
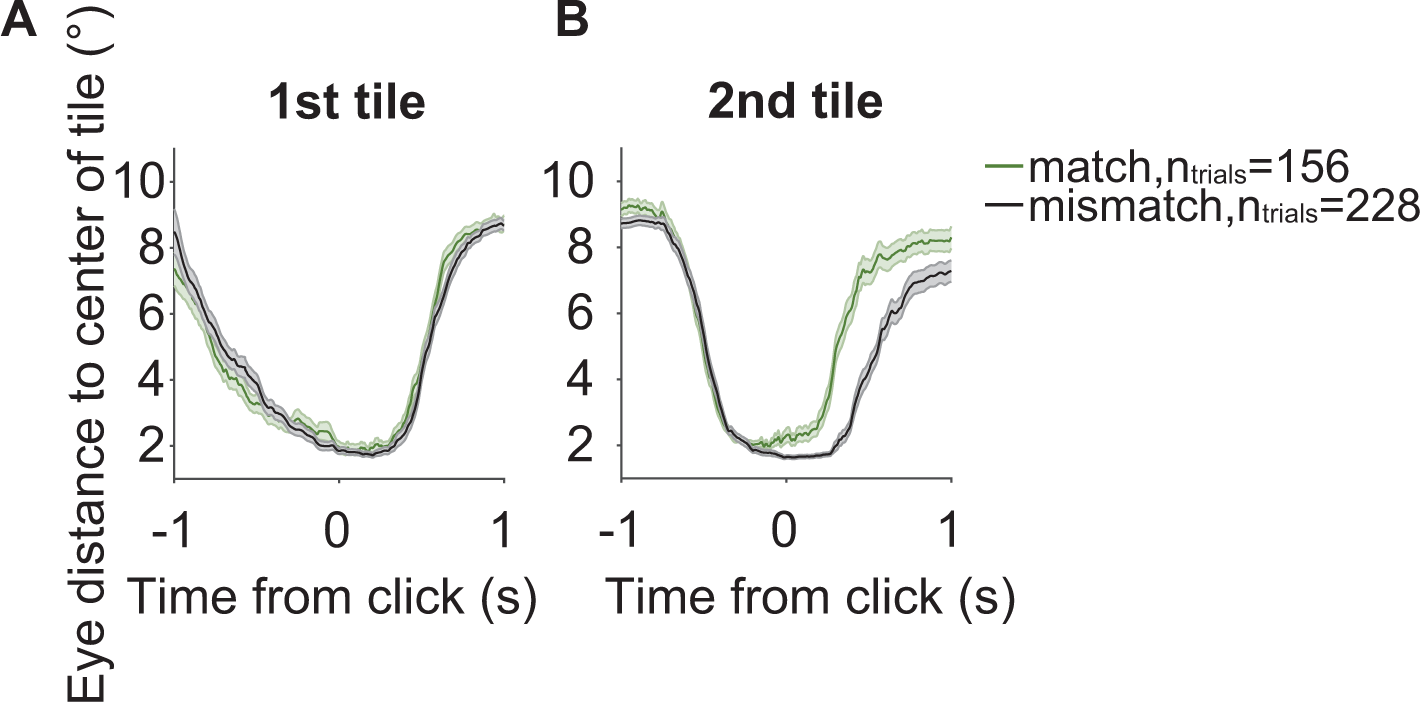
Participants’ eye movements tracked the clicked locations. Eye movements were similar during match and mismatch trials for the 1^st^ tile, and for the 2^nd^ tile before click time, but not for the 2^nd^ tile after click time, when equalizing the reaction time and distances between the 1^st^ and 2^nd^ tiles. Line traces (mean±s.e.m) indicate the average distance in visual angle between gaze and the center of the tile clicked for match (green) and mismatch (black) trials. Reaction times (RT) and distances between the 1^st^ and 2^nd^ tiles were equalized (mean RT ± 200ms; mean distance ± 3
°). **A**. No significant difference in eye-movements between match and mismatch was found during clicking the 1^st^ tile. **B**. No significant difference in eye movements between match and mismatch was found before clicking the 2^nd^ tile. Participants’ gaze moved faster away from the 2^nd^ tile in match trials than mismatch trials after the click: within the 1 second window after time zero, the distance from the gaze to the center of the tile was on average 1.76 dva (degrees of visual angle) larger for match than mismatch trials.

**Figure S2.**
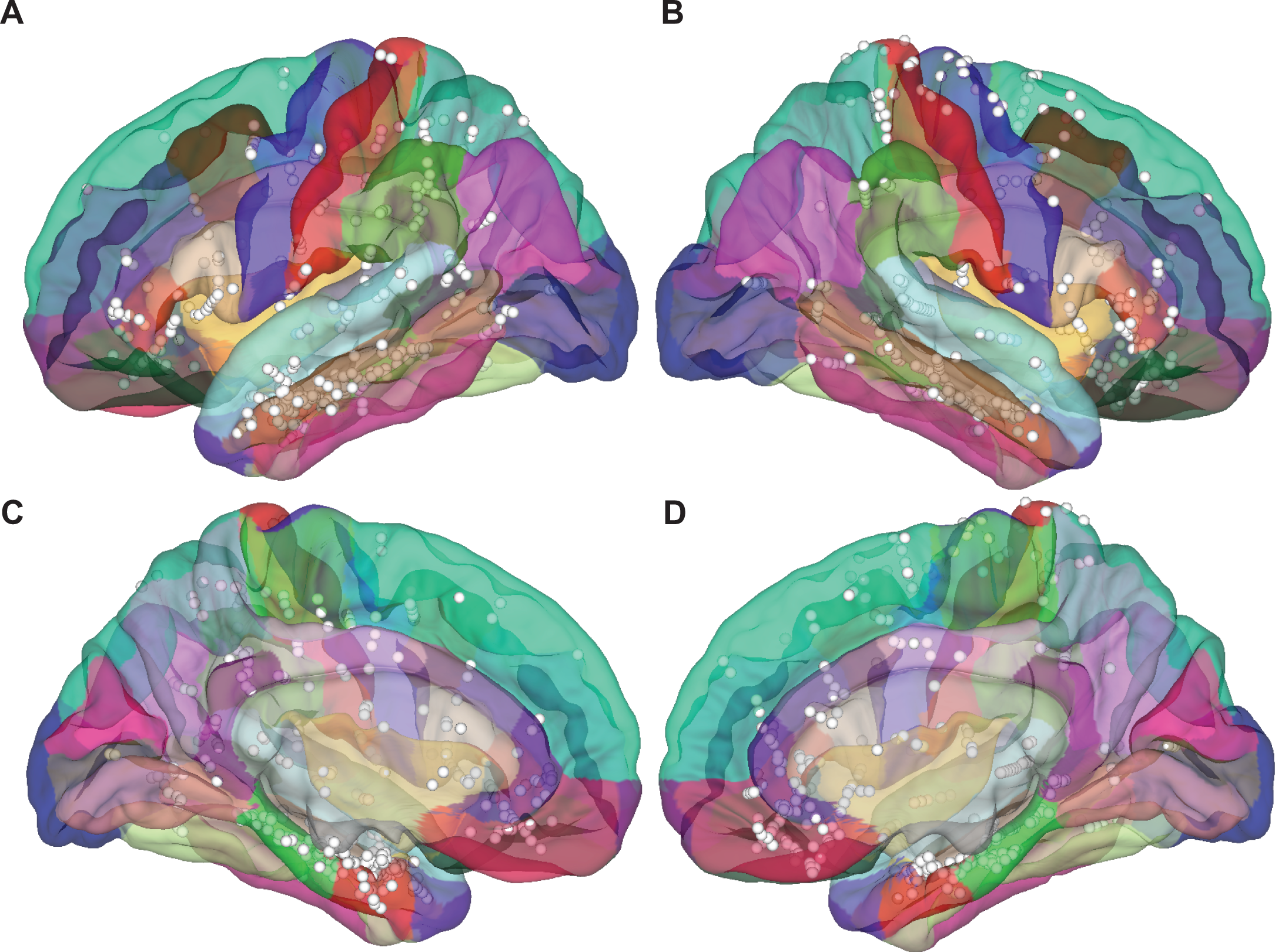
Locations of electrodes in the gray matter Each circle shows a bipolarly referenced electrode (n=676), overlayed on the Desikan-Killiany Atlas with different views: **A**: left lateral; **B**: right lateral; **C**: left medial; **D**: right medial. The colors reflect the Desikan-Killiany parcellation (Desikan et al, 2006). Locations of white matter electrodes are reported in **Figure S3** (see also **Table S2**).

**Figure S3.**
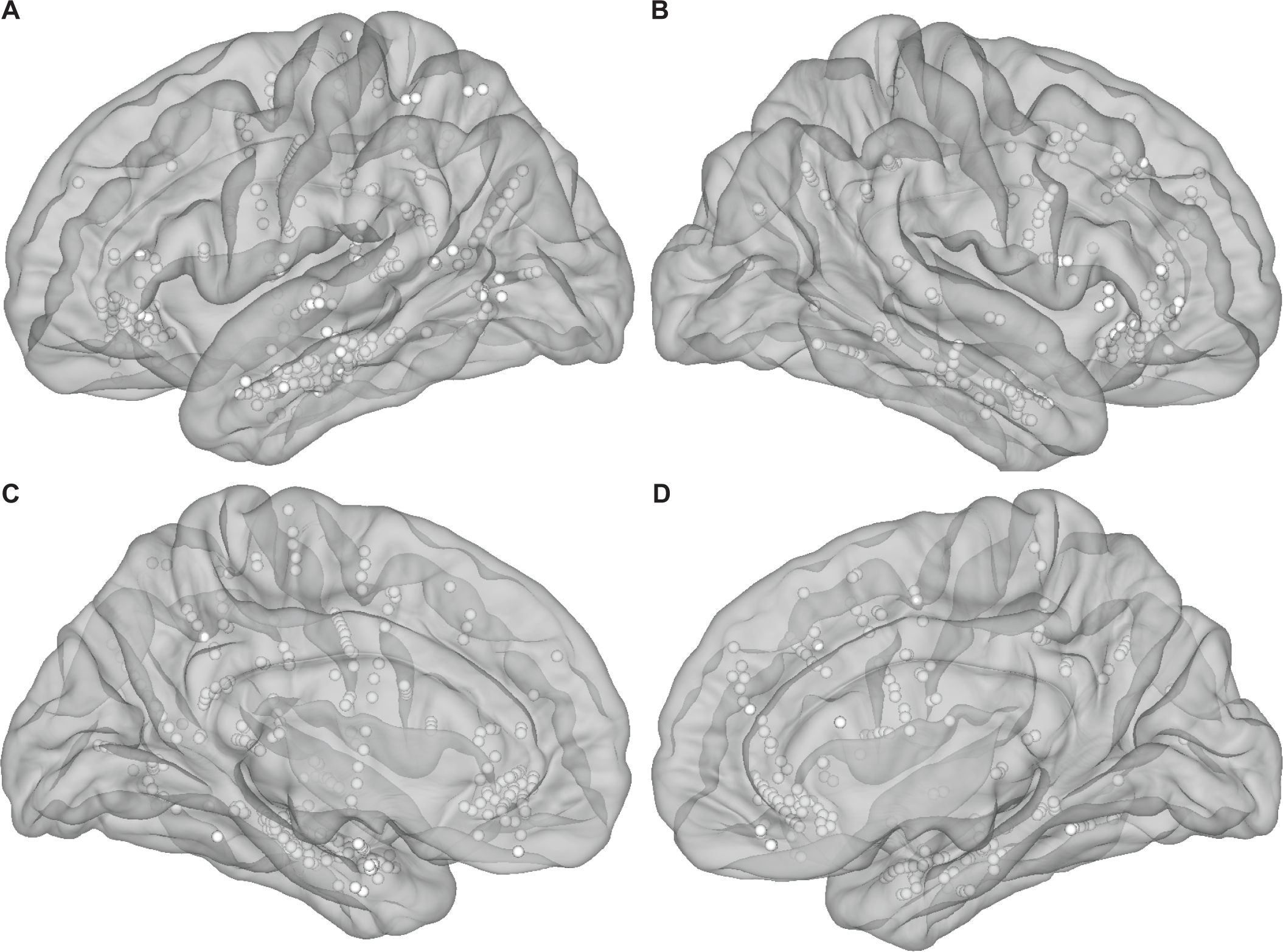
Locations of all analyzed electrodes in the white matter. Each sphere reflects one of each pair of nearby electrodes that were bipolarly referenced (n=492, **Table S2**). **A**: left lateral view; **B**: right lateral view; **C**: left medial view; **D**: right medial view. The location of gray matter electrodes is shown in **Figure S2**.

**Figure S4.**
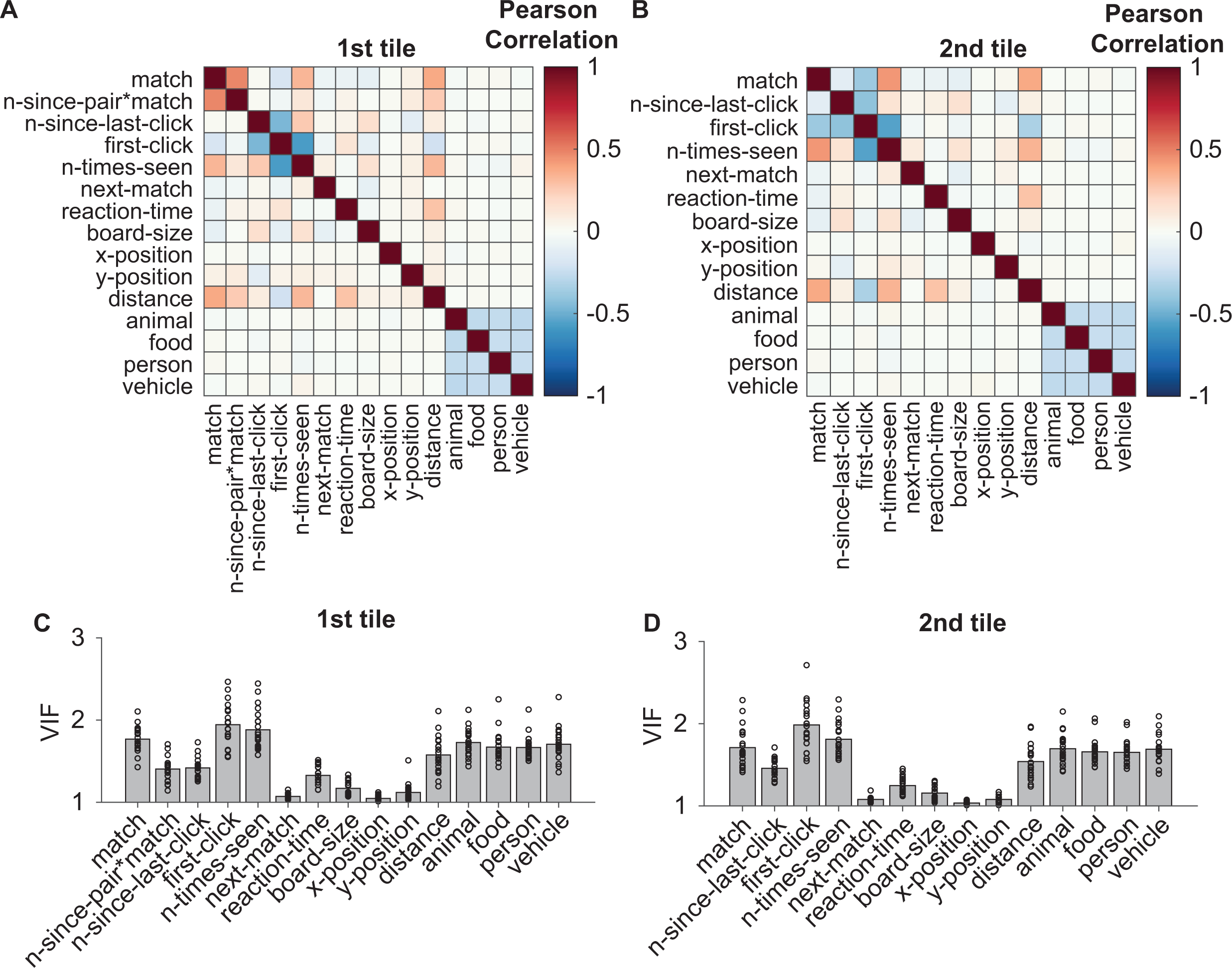
Correlation and collinearity among predictor variables**. A**, **B**. Average across subjects of the Pearson correlation between pairs of predictors for the 1^st^ tile (**A**) and the 2^nd^ (**B**) tile. See color scale on the right. This is a symmetric matrix and the diagonal is 1, by definition. Table 1 describes each of the predictors. **C**, **D**. Variation Inflation Factor (VIF) (O’Brien, 2007) of each predictor for the 1^st^ tile (**C**) and the 2^nd^ (**D**) tile. Bar height indicates the average VIF of each predictor across all subjects. Dots represent the VIF of each predictor for each subject.

**Figure S5.**
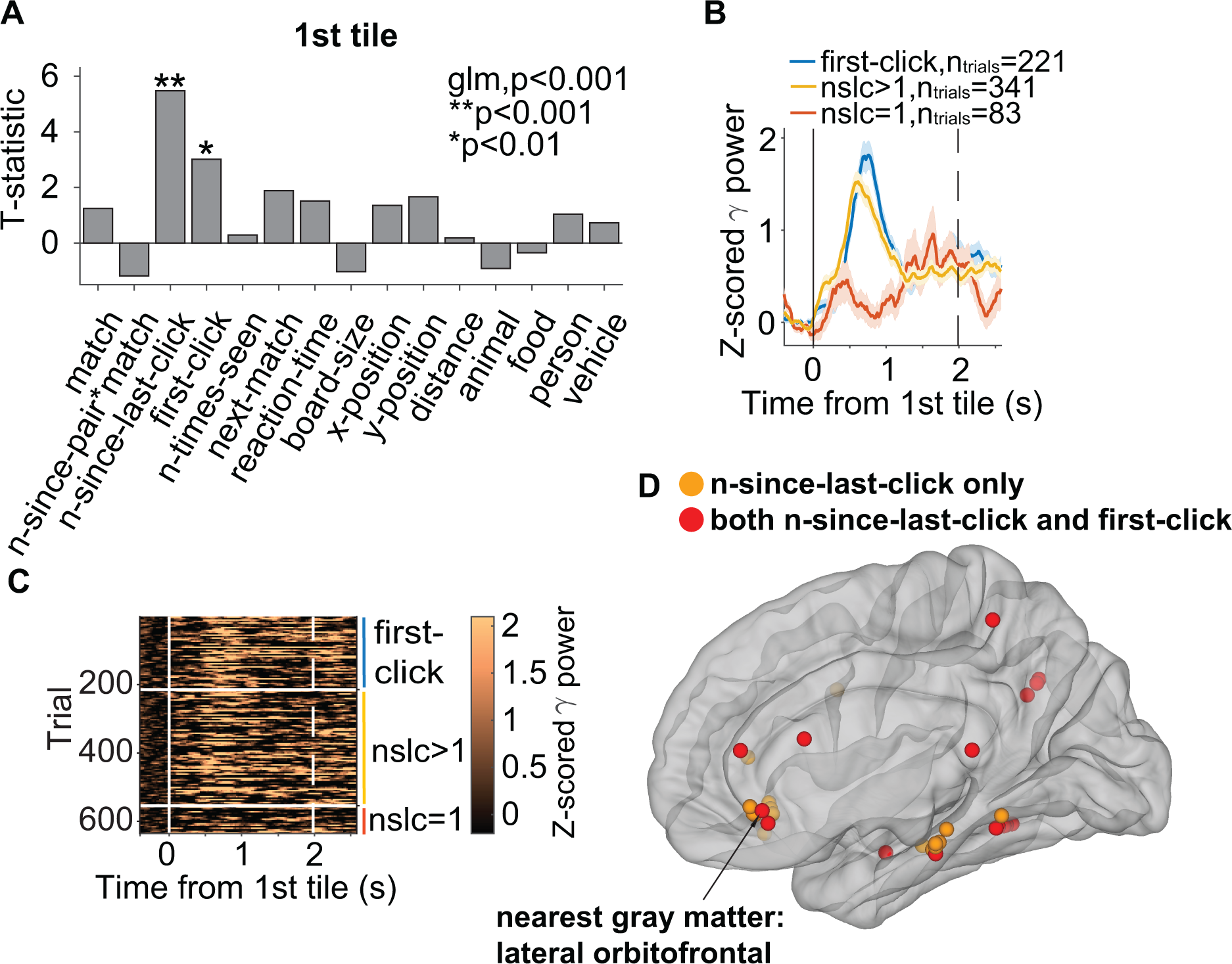
Neural signals in the white matter reflect novelty and familiarity Panels show an example electrode in the white matter whose closest gray matter region is the left lateral orbitofrontal cortex (arrow in **D**). This figure follows the format of **Figure 3** in the main text. **A.** T-statistic of each predictor in the GLM analysis (**Methods**). Asterisks indicate statistically significant predictors for the neural signals. **B.** Z-scored gamma band power aligned to the 1^st^ tile onset (solid vertical line) for novel tiles (blue), unfamiliar tiles (n-since-last-click>1, yellow), and more familiar tiles (n-since-last-click=1, red). The vertical dashed line indicates the mean reaction time. The time axis extends from 400 ms before the click to 500 ms after the average reaction time. **C.** Raster plot showing the z-scored gamma power in individual trials ordered by first-click and then larger to smaller n-since-last-click; division indicated by white horizontal lines and colored vertical lines. **D.** Locations of all white matter electrodes where n-since-last-click was a significant predictor during the 1st tile. Orange: n-since-last-click; red: both n-since-last-click and first-click were significant predictors. All electrodes were reflected on one hemisphere for display purposes.

**Figure S6.**
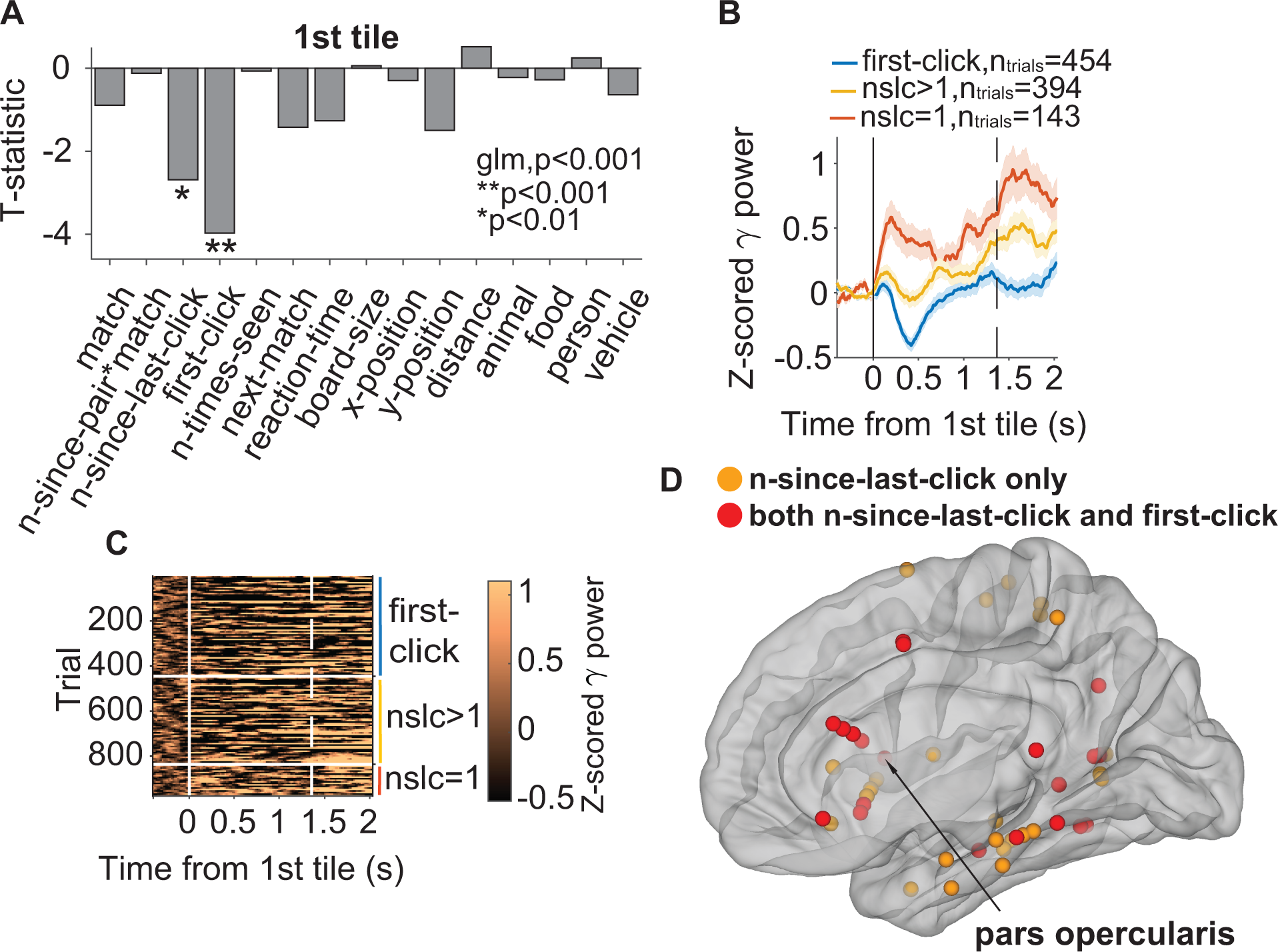
Neural signals reflect novelty and familiarity with a pattern different from that shown in Figure 3E-G. This figure shows an example electrode in the right pars opercularis (gray matter, arrow in **D**), following the format of **Figure 3** in the main text. In contrast to **Figure 3E-G**, this electrode showed increased activity for more familiar trials. **A.** T-statistic of each predictor in the GLM for the 1st tile. Asterisks indicate significant predictors for the gamma power AUC. **B.** Z-scored gamma (30-150 Hz) power for first-click (blue), n-since-last-click>1 (yellow), and n-since-last-click=1 (red). Dashed line indicates the mean reaction time. **C.** Raster plot showing the z-scored gamma power in individual trials ordered by first-click and then larger to smaller n-since-last-click; division indicated by white horizontal lines and colored vertical lines. **D.** Locations of all n-since-last-click electrodes during the 1st tile. Orange: n-since-last-click only; red: both n-since-last-click and first-click were significant predictors. All electrodes were reflected on one hemisphere for display purposes.

**Figure S7.**
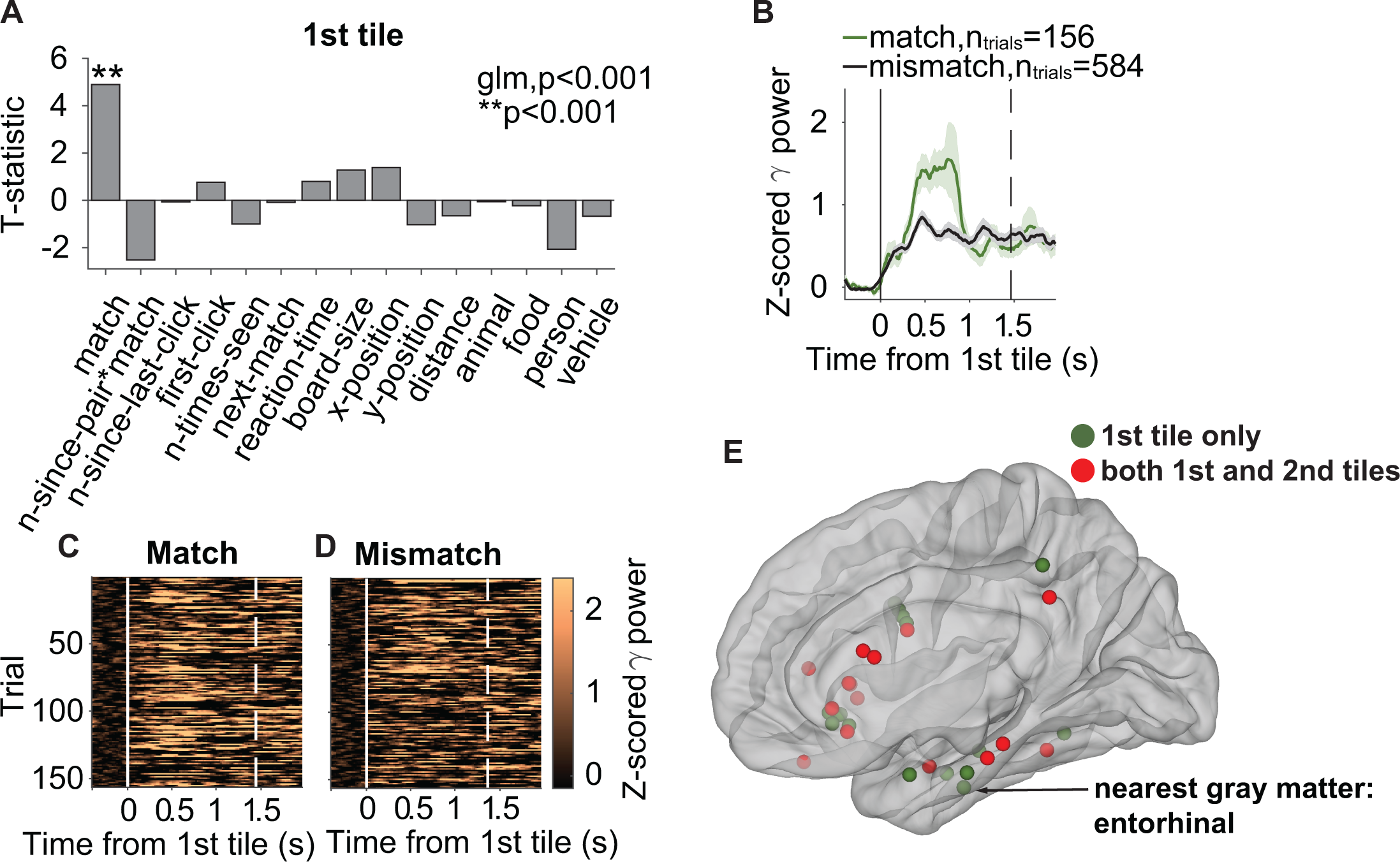
Neural signals in the white matter predict correct retrieval Panels show an example electrode located in the white matter where the nearest gray matter region is the right entorhinal cortex (see arrow in **E**) as well as population locations in **E**. The format follows **Figure 3**. **A.** t-statistic of each predictor in the GLM analysis. Asterisks indicate statistically significant predictors for the neural signals. **B.** Z-scored gamma band power aligned to the 1^st^ tile onset (solid vertical line) for match trials (green) and mismatch trials (black). The vertical dashed line indicates the mean reaction time. Shaded error bars indicate s.e.m. **C**-**D**. Raster plots showing the gamma power in individual trials for match (**C**) and mismatch (**D**) trials. **E.** Locations of all the white matter electrodes where match was a significant predictor during the 1st tile only (green) and during both tiles (red). All electrodes were reflected on one hemisphere for display purposes.

**Figure S8.**
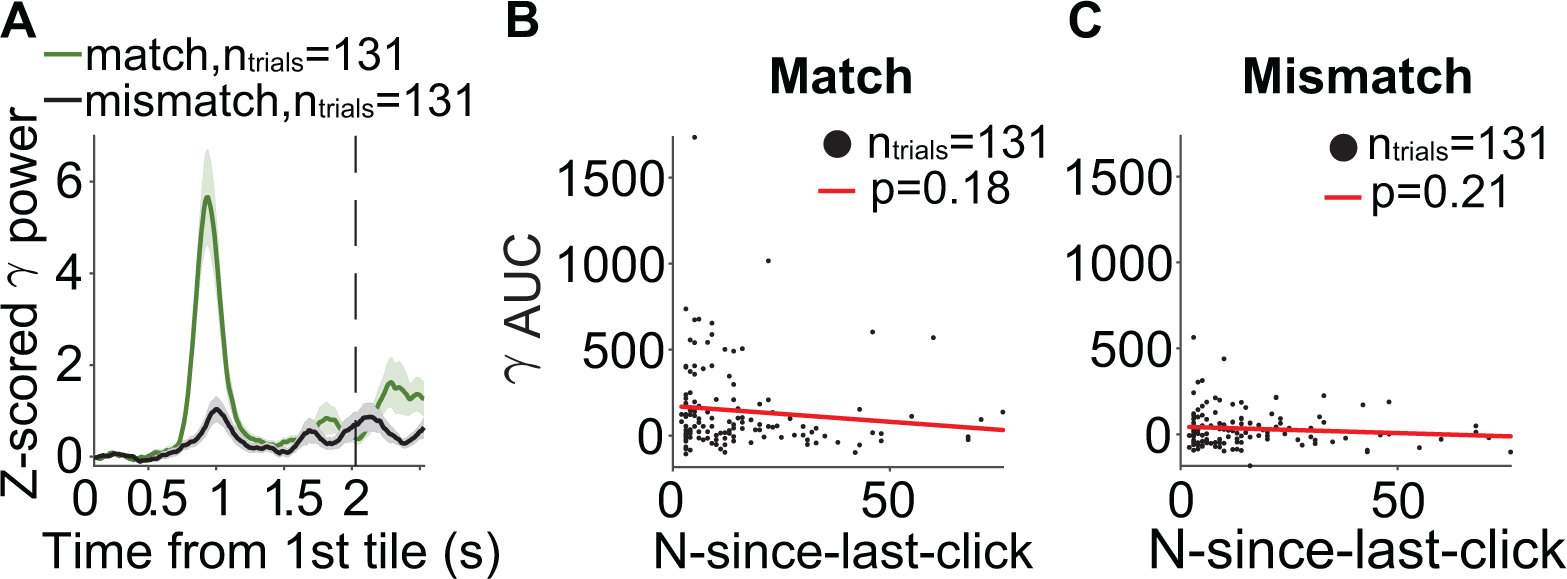
Match is the only predictor that accounts for the neural differences between match and mismatch Panels shown the neural responses of an example electrode in the right lateral orbitofrontal cortex (same as in **Figure 4**) when equalizing the distribution of n-since-last click in match and mismatch trials. **A.** Z-scored gamma (30-150 Hz) power during match (green) and mismatch (black) trials aligned to the 1st tile onset (solid line) after equalizing n-since-last-click values of match and mismatch trials by random subsampling. Dashed line indicates the mean reaction time. **B-C.** Scatter plots of AUC of gamma power versus n-since-last-click for match (**B**) and mismatch (**C**) trials. Each dot represents data from one trial. Red lines represent linear fits of the data.

**Figure S9.**
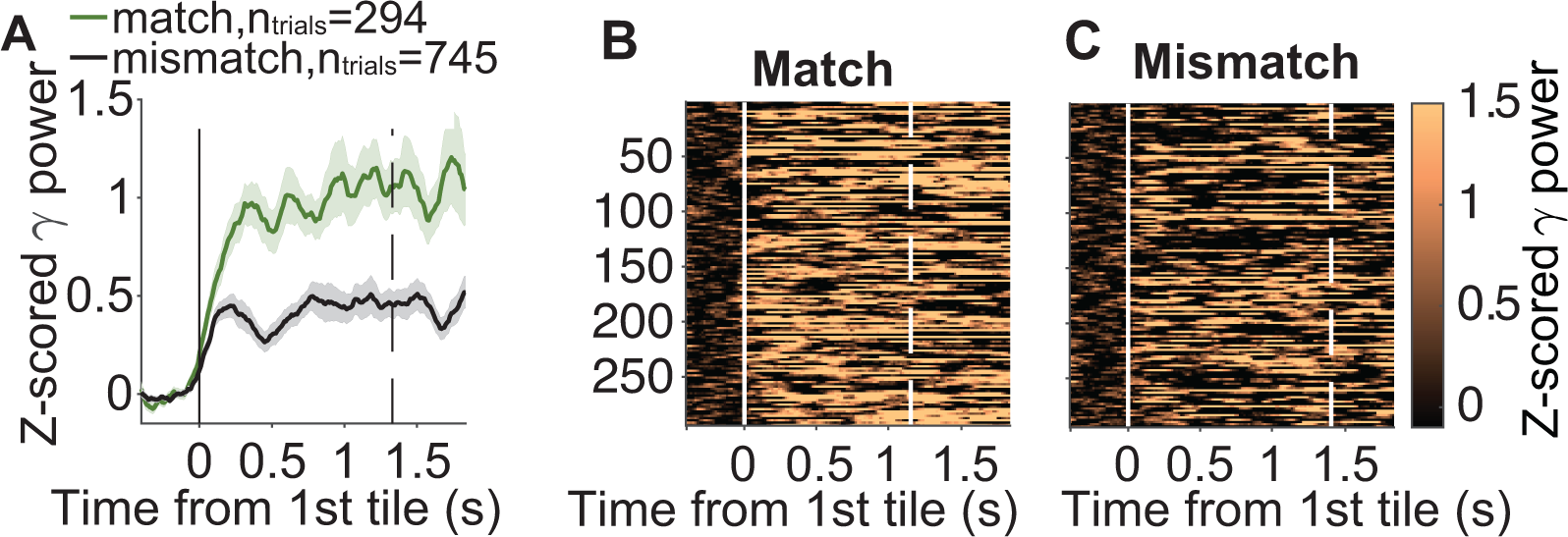
Neural signals predict correct retrieval with a pattern different from that shown in Figure 4 Panels show an example electrode located in the left middle temporal gyrus where “match” was a significant predictor for gamma activity. **A**. Z-scored gamma (30-150 Hz) power during match (green) and mismatch (black) trials aligned to the 1st tile onset (solid line). Dashed line indicates the mean reaction time. **B**-**C**. Raster plots showing the z-scored gamma power in individual match (**B**) and mismatch (C) trials. For display purposes, trial numbers of match and mismatch were equalized (**Methods**).

**Figure S10.**
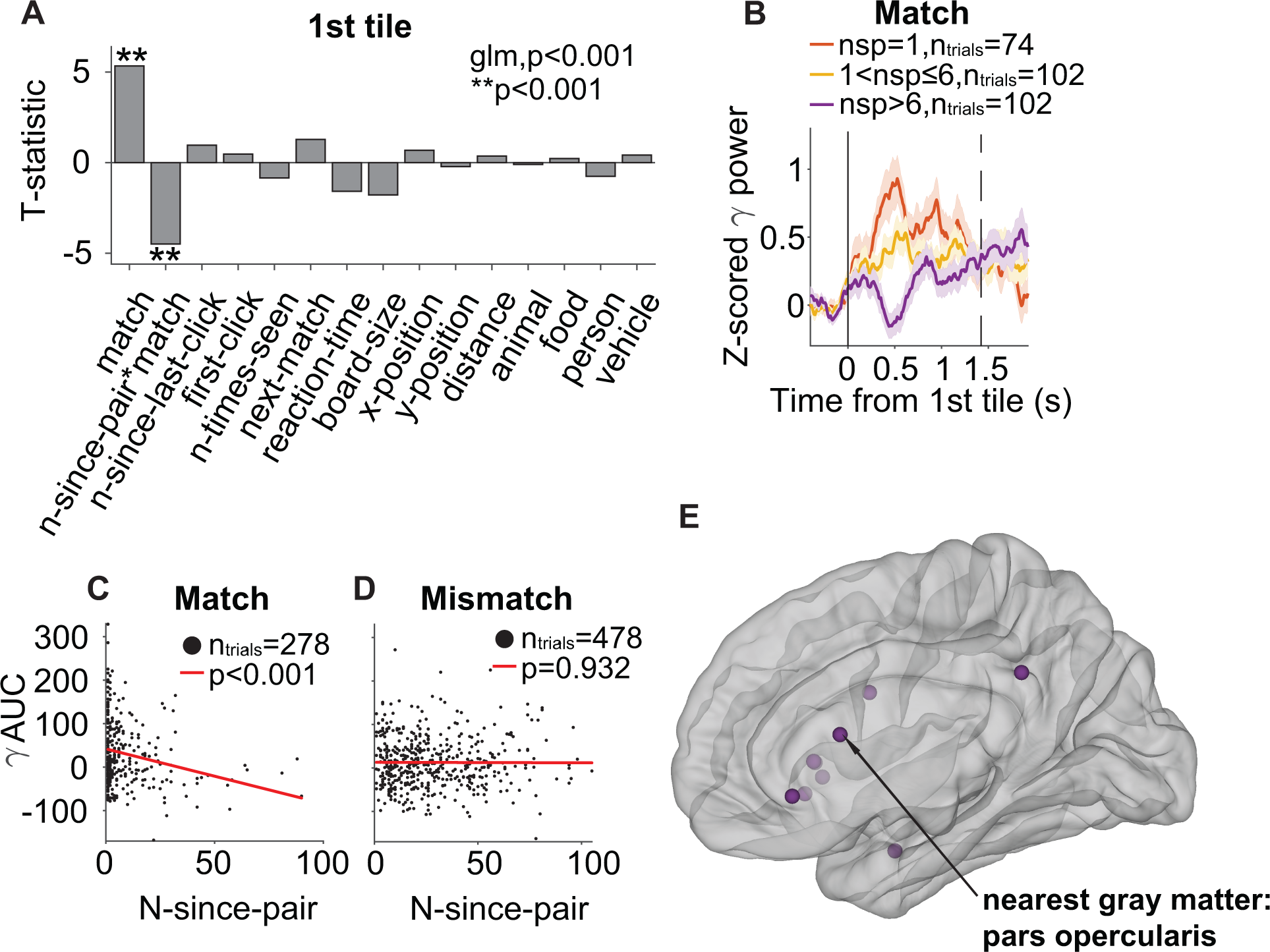
Neural signals in the white matter reflect the strength of memory retrieval. Panels show an example electrode in the white matter where the closest gray matter region is the left pars opercularis (see arrow in part **E**) and population locations in **E**. **A.** T-statistic of each predictor in the GLM analysis. Asterisks indicate significant predictors for the neural signals. **B.** Z-scored gamma band power aligned to the 1^st^ tile onset (solid vertical line) for match trials with small n-since-pair (red, stronger memories), intermediate nsp (yellow), and large nsp (purple, weaker memories). The vertical dashed line indicates the mean reaction time. Shaded error bars indicate s.e.m. **C-D**. Scatter plots of the area under the curve (AUC) of the gamma band power as a function of nsp for match trials (**C**) and mismatch trials (**D**). Each dot represents one trial. Red lines show linear fits to the data. **B.** Locations of all white matter electrodes where nsp was a significant predictor. All electrodes were reflected on one hemisphere for display purposes.

**Figure S11.**
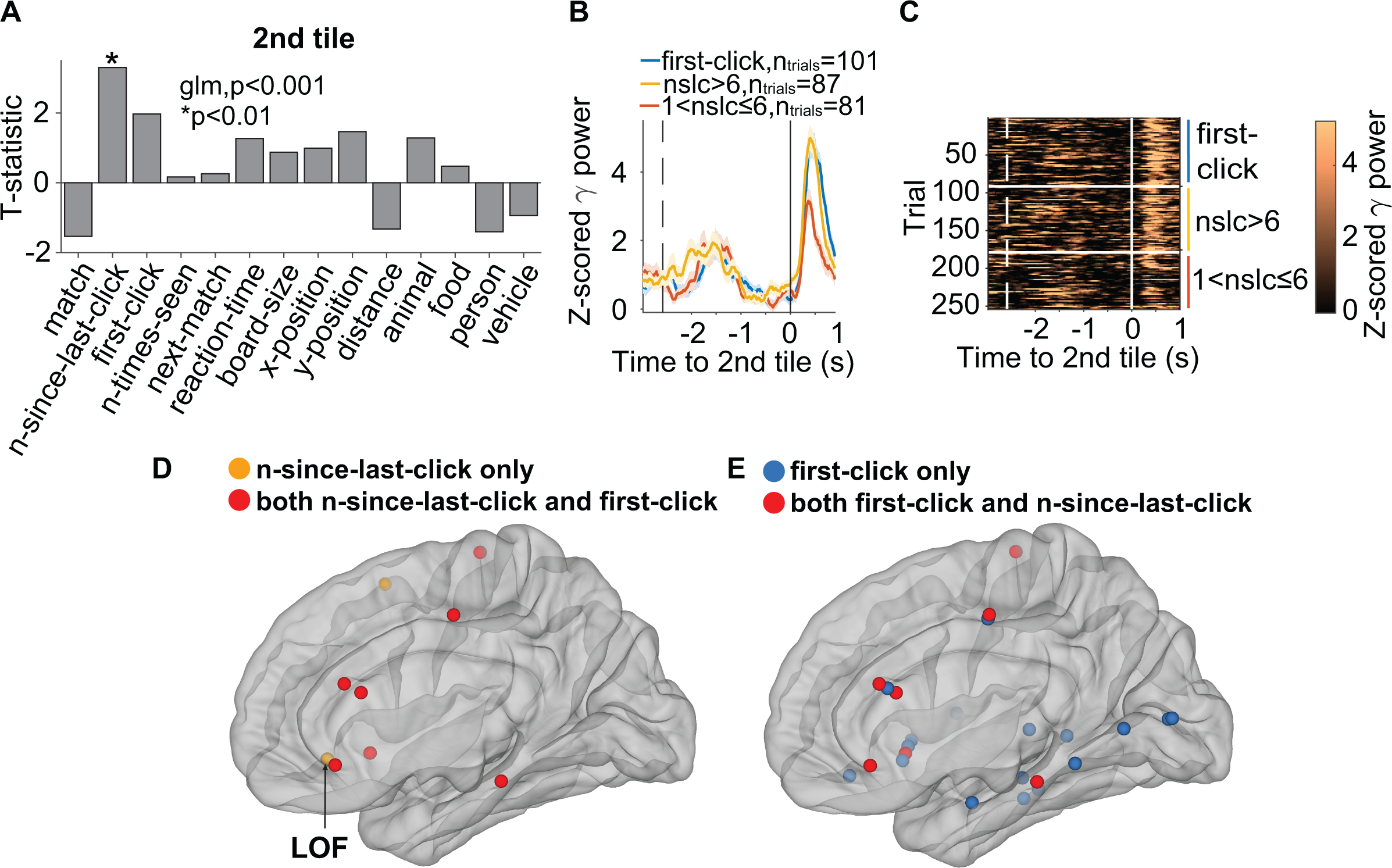
Neural signals after the second tile reflect familiarity and novelty Panels show an example electrode in the right lateral orbitofrontal (LOF) cortex and population locations in **D**-**E**. **A.** T-statistic of each predictor in the GLM analysis (**Methods**). Asterisks indicate statistically significant predictors for the neural signals. **B.** Z-scored gamma band power aligned to the 2^nd^ tile onset (solid vertical line) for novel tiles (blue), unfamiliar tiles (n-since-last-click>6, yellow), and familiar tiles (n-since-last-click≤1, red). The vertical dashed line indicates the mean onset of the 1st tile. The time axis extends from 400 ms plus the average reaction time before the click to 1,000 ms after the click. **C.** Raster plots showing the z-scored gamma power in individual trials ordered by first-click and then larger to smaller n-since-last-click; division indicated by white horizontal lines/spaces and colored vertical bars. **D.** Locations of all electrodes where n-since-last-click was a significant predictor during the 2nd tile. Orange: n-since-last-click only; red: both n-since-last-click and first-click were significant predictors. **E.** Locations of all electrodes where first-click was a significant predictor during the 2nd tile. Blue: first-click only; red: both first-click and n-since-last-click were significant predictors. All electrodes were reflected on one hemisphere for display purposes.

**Figure S12.**
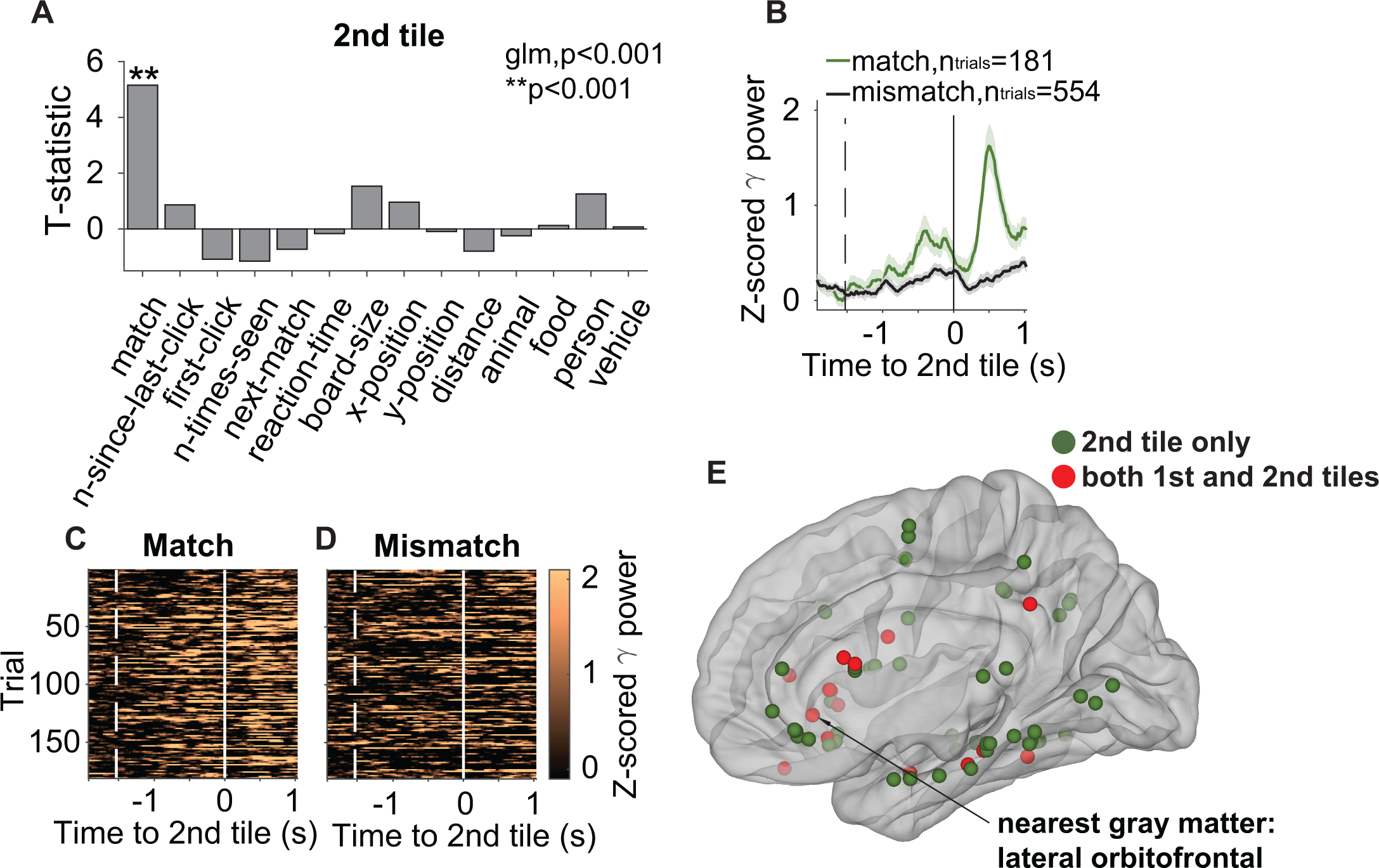
Neural signals in the white matter after the second tile reflect correct retrieval Panels show an example electrode located in the white matter whose closest gray matter region is the right LOF (see arrow in part **E**). **A**. T-statistic of each predictor in the GLM analysis for the responses after the 2^nd^ tile. Asterisks indicate statistically significant predictors of neural signals. **B**. Z-scored gamma band power aligned to the onset of the 2^nd^ tile (solid vertical line) for match (green) and mismatch (black) trials. The dashed line indicates the mean onset of the 1^st^ tile. **C**-**D**. Raster plots showing the gamma band power in individual trials for match (left) and mismatch (right) trials. **E**. Locations of all the white matter electrodes where match was a significant predictor during the 2nd tile only (green) and during both tiles (red). All electrodes were reflected on one hemisphere for display purposes.

**Figure S13.**
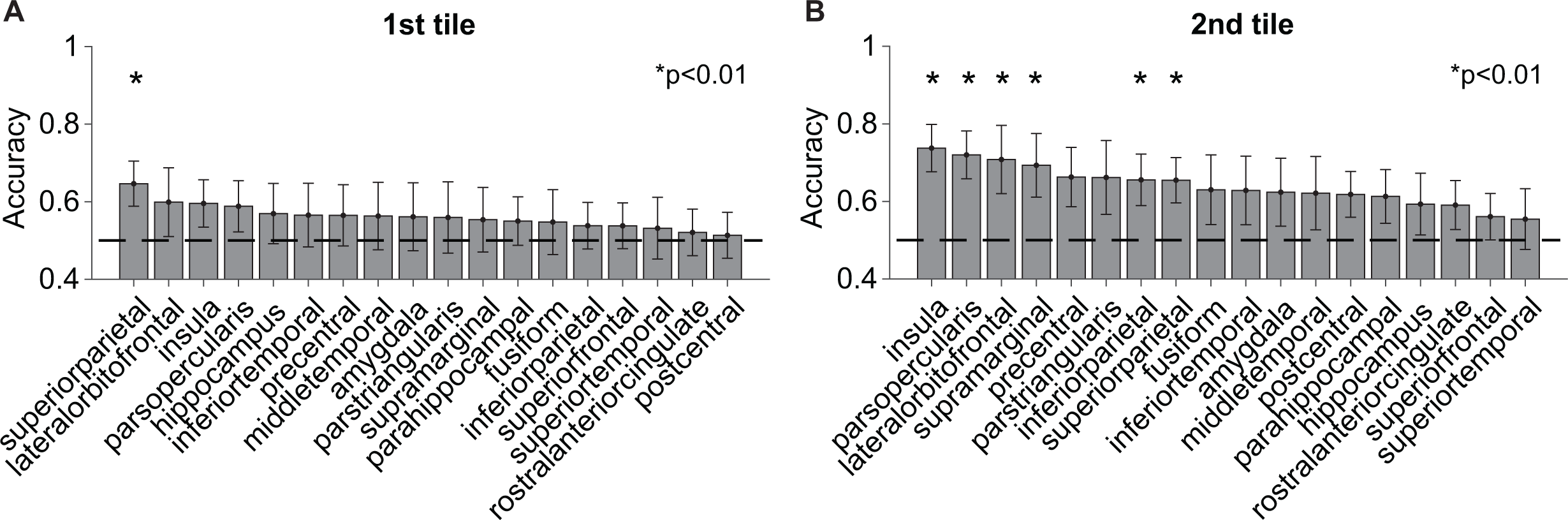
Decoding of match versus mismatch with random subsampling of 12 channels in each brain region. Average decoding accuracy of each brain region (**Methods**) using neural responses after the 1^st^ tile (**A**) and the 2^nd^ tile (**B)**. The format follows **Figure 7**. Brain regions are ordered from higher to lower average decoding accuracy. Dashed horizontal lines indicate chance accuracy. Asterisks denote significant decoding accuracy above chance (⍺=0.01). All error bars indicate SD (n=1,000 iterations).

**Table S1:**
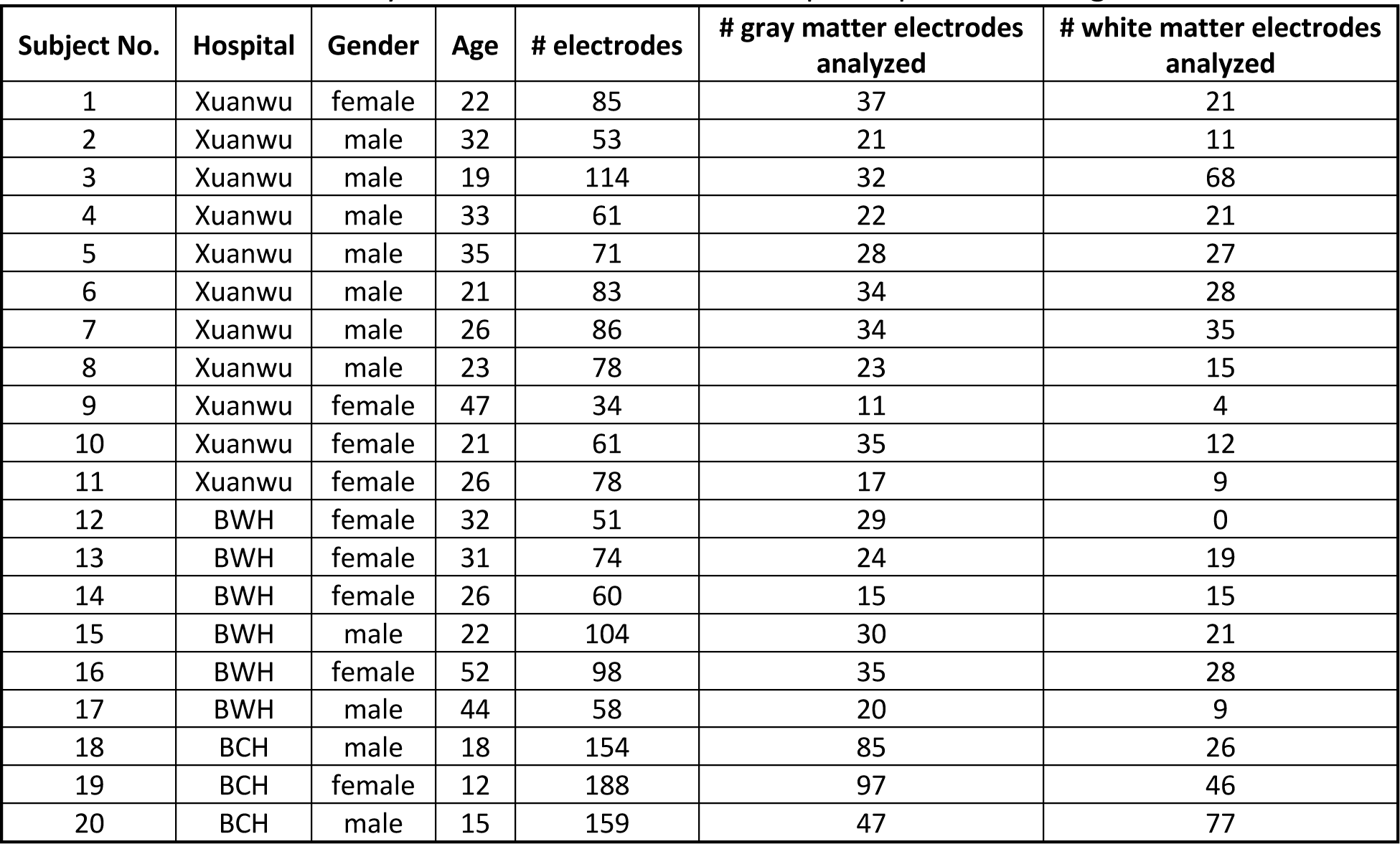
Information about each participant: hospital where data were recorded, gender, age, total number of electrodes implanted, number of electrodes analyzed in the gray matter, and number of electrodes analyzed in the white matter. All participants were right-handed.

**Table S2:** Distribution of electrode locations per hemisphere and brain region for the gray matter (left) and white matter (right). The location of any white matter electrode was based on its nearest gray matter region.

| Gray matter |  |  |  | White matter (closest gray matter) |  |  |  |
| --- | --- | --- | --- | --- | --- | --- | --- |
| Region | Left | Right | Total | Region | Left | Right | Total |
| amygdala | 21 | 24 | 45 | amygdala | 0 | 0 | 0 |
| bankssts | 0 | 7 | 7 | bankssts | 5 | 2 | 7 |
| caudalanteriorcingulate | 1 | 3 | 4 | caudalanteriorcingulate | 1 | 0 | 1 |
| caudalmiddlefrontal | 1 | 5 | 6 | caudalmiddlefrontal | 1 | 7 | 8 |
| entorhinal | 5 | 0 | 5 | cuneus | 1 | 1 | 2 |
| fusiform | 15 | 10 | 25 | entorhinal | 2 | 2 | 4 |
| hippocampus | 36 | 31 | 67 | fusiform | 21 | 14 | 35 |
| inferiorparietal | 11 | 8 | 19 | inferiorparietal | 7 | 3 | 10 |
| inferiortemporal | 14 | 9 | 23 | inferiortemporal | 31 | 16 | 47 |
| insula | 13 | 15 | 28 | insula | 11 | 12 | 23 |
| isthmuscingulate | 6 | 0 | 6 | isthmuscingulate | 1 | 0 | 1 |
| lateraloccipital | 3 | 2 | 5 | lateraloccipital | 2 | 3 | 5 |
| lateralorbitofrontal | 19 | 32 | 51 | lateralorbitofrontal | 23 | 30 | 53 |
| lingual | 3 | 2 | 5 | lingual | 2 | 2 | 4 |
| medialorbitofrontal | 4 | 7 | 11 | medialorbitofrontal | 6 | 8 | 14 |
| middletemporal | 53 | 38 | 91 | middletemporal | 31 | 21 | 52 |
| paracentral | 4 | 0 | 4 | paracentral | 3 | 3 | 6 |
| parahippocampal | 10 | 13 | 23 | parahippocampal | 6 | 5 | 11 |
| parsopercularis | 11 | 5 | 16 | parsopercularis | 5 | 3 | 8 |
| parsorbitalis | 2 | 3 | 5 | parsorbitalis | 4 | 7 | 11 |
| parstriangularis | 13 | 19 | 32 | parstriangularis | 10 | 10 | 20 |
| pericalcarine | 0 | 1 | 1 | pericalcarine | 4 | 0 | 4 |
| postcentral | 8 | 12 | 20 | postcentral | 6 | 0 | 6 |
| posteriorcingulate | 4 | 3 | 7 | posteriorcingulate | 10 | 1 | 11 |
| precentral | 8 | 21 | 29 | precentral | 11 | 15 | 26 |
| precuneus | 3 | 0 | 3 | precuneus | 12 | 3 | 15 |
| rostralanteriorcingulate | 7 | 7 | 14 | rostralanteriorcingulate | 3 | 3 | 6 |
| rostralmiddlefrontal | 1 | 4 | 5 | rostralmiddlefrontal | 2 | 7 | 9 |
| superiorfrontal | 13 | 19 | 32 | superiorfrontal | 10 | 18 | 28 |
| superiorparietal | 8 | 12 | 20 | superiorparietal | 10 | 7 | 17 |
| superiortemporal | 26 | 21 | 47 | superiortemporal | 21 | 13 | 34 |
| supramarginal | 10 | 4 | 14 | supramarginal | 4 | 4 | 8 |
| transversetemporal | 6 | 0 | 6 | transversetemporal | 6 | 0 | 6 |
| **Total** | **339** | **337** | **676** | **Total** | **272** | **220** | **492** |

**Table S3A.**
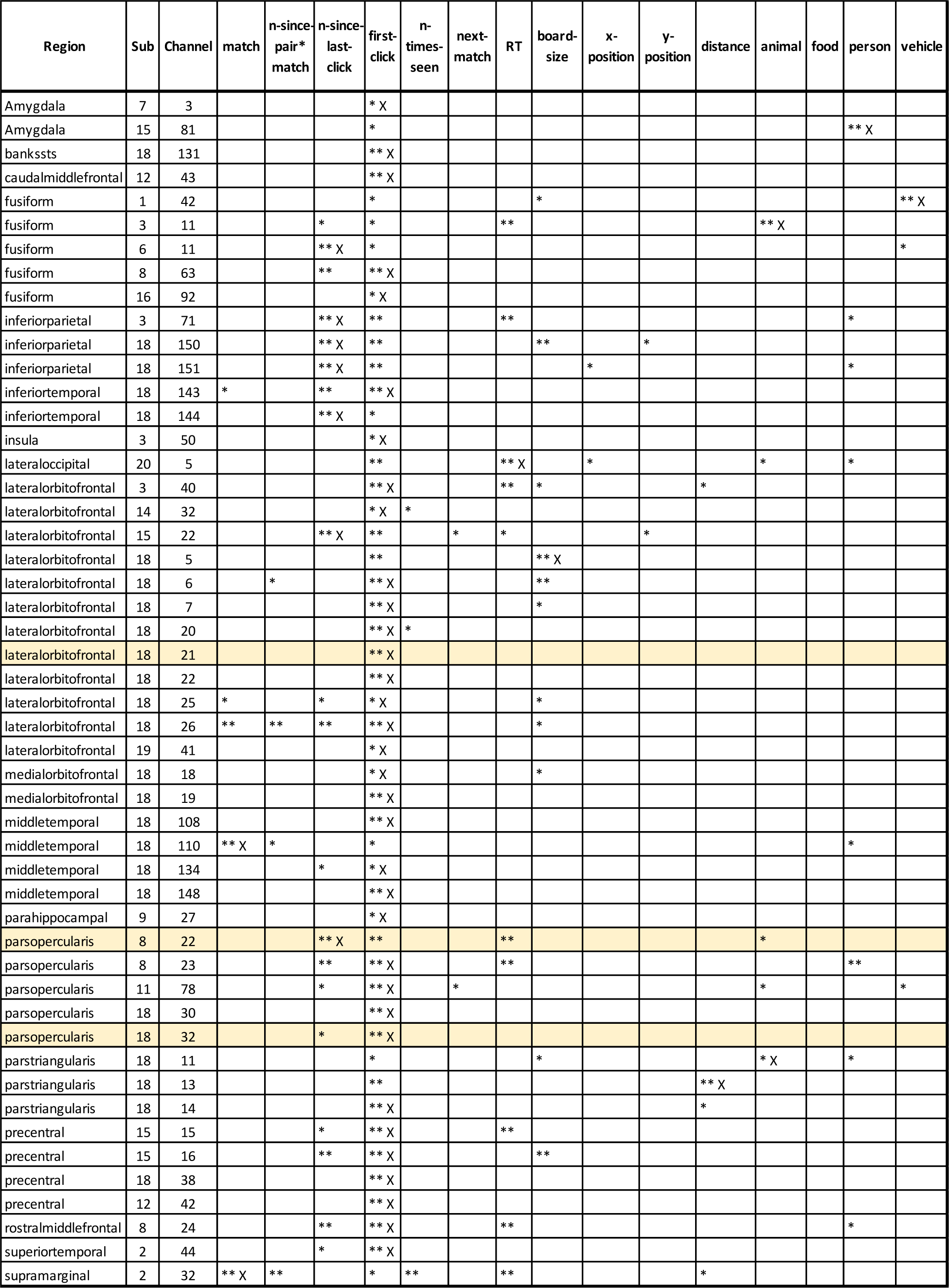
First tile GLM analysis for all gray matter electrodes that showed first-click as a significant predictor. Each column (starting from column 4) shows a separate predictor. Yellow rows indicate the example electrodes in **Figure 4** (first and second rows) and **Figure S4** (third row). *: p<0.01. **: p<0.001. X: most significant predictor.

**Table S3B.**
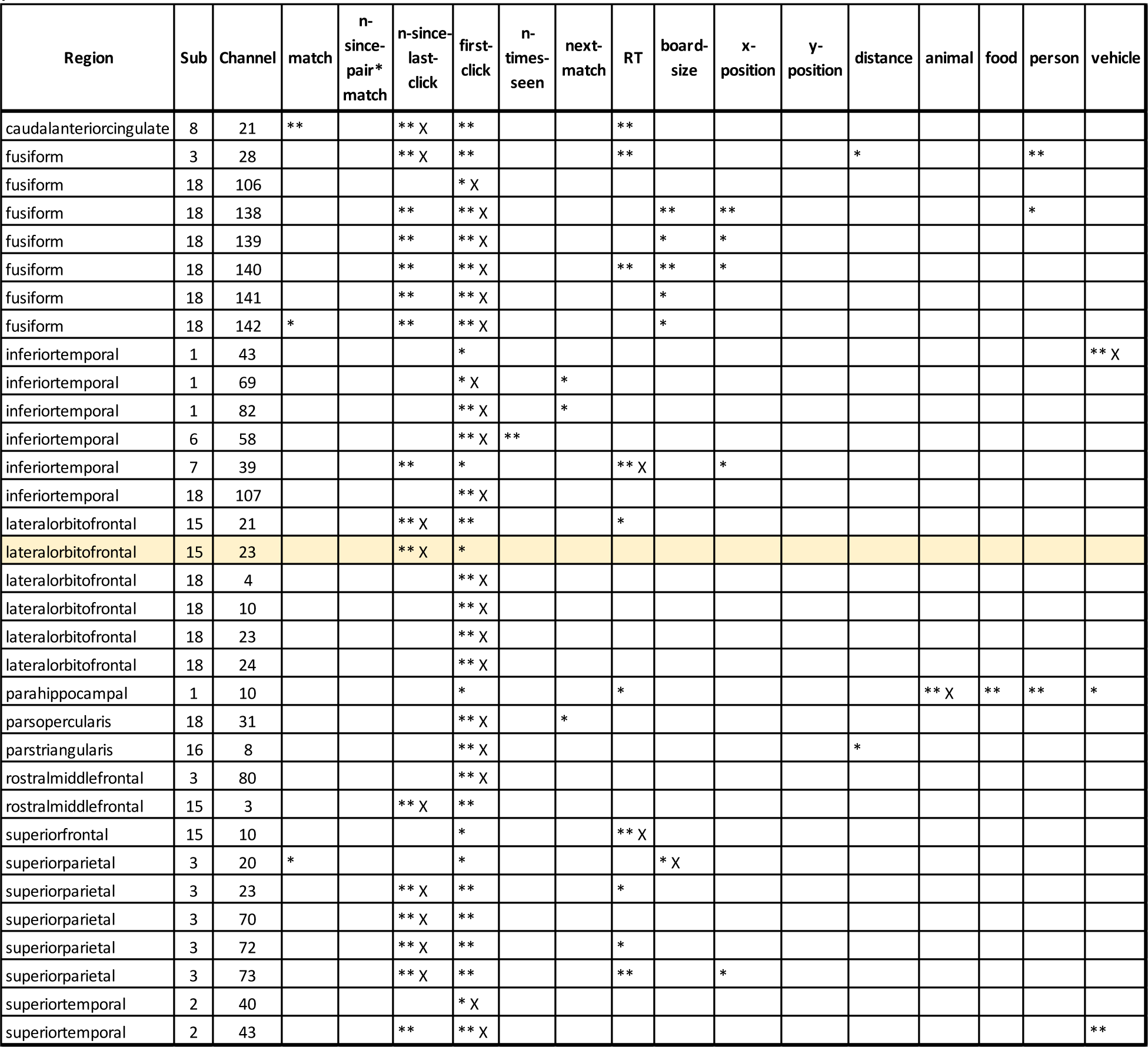
First tile GLM analysis for all white matter electrodes that showed first-click as a significant predictor. Each column (starting from column 4) shows a separate predictor. Yellow row indicates the example electrode in **Figure S4**. *: p<0.01. **: p<0.001. X: most significant predictor.

**Table S4A.** First tile GLM analysis for all gray matter electrodes that showed n-since-last-click as a significant predictor. Each column (starting from column 4) shows a separate predictor. Yellow rows indicate the example electrodes in **Figure 4E-H** (first row) and **Figure S5** (second row).*: p<0.01. **: p<0.001. X: most significant predictor.

| Region | Sub | Channel | match | n-since-pair*<br>match | n-since-last-click | first-click | n-times-seen | next-match | RT | board-size | x-position | y-position | distance | animal | food | person | vehicle |
| --- | --- | --- | --- | --- | --- | --- | --- | --- | --- | --- | --- | --- | --- | --- | --- | --- | --- |
| Hippocampus | 6 | 6 |  |  | * |  | * X |  |  |  |  |  |  |  |  |  |  |
| bankssts | 16 | 41 |  |  | * |  |  |  |  |  |  |  |  | * |  | ** X |  |
| bankssts | 16 | 42 |  |  | * |  |  |  |  |  |  |  |  | * | * | ** X |  |
| entorhinal | 6 | 27 | * |  | * X |  |  |  |  |  |  |  |  |  |  |  |  |
| fusiform | 2 | 13 |  |  | * X |  |  |  |  |  |  |  |  |  |  |  |  |
| fusiform | 3 | 11 |  |  | * | * |  |  | ** |  |  |  |  | ** X |  |  |  |
| fusiform | 6 | 11 |  |  | ** X | * |  |  |  |  |  |  |  |  |  |  | * |
| fusiform | 8 | 63 |  |  | ** | ** X |  |  |  |  |  |  |  |  |  |  |  |
| inferiorparietal | 3 | 71 |  |  | ** X | ** |  |  | ** |  |  |  |  |  |  | * |  |
| inferiorparietal | 18 | 150 |  |  | ** X | ** |  |  |  | ** |  | * |  |  |  |  |  |
| inferiorparietal | 18 | 151 |  |  | ** X | ** |  |  |  |  | * |  |  |  |  | * |  |
| inferiorparietal | 18 | 152 |  |  | ** X |  |  |  |  |  |  |  |  |  |  |  |  |
| inferiortemporal | 7 | 38 | * |  | * |  |  |  | ** X |  | * |  |  |  |  |  |  |
| inferiortemporal | 18 | 143 | * |  | ** | ** X |  |  |  |  |  |  |  |  |  |  |  |
| inferiortemporal | 18 | 144 |  |  | ** X | * |  |  |  |  |  |  |  |  |  |  |  |
| insula | 18 | 27 | ** | ** | ** X |  |  |  |  |  |  |  |  |  |  |  |  |
| insula | 18 | 28 | ** | ** | ** X |  |  |  |  |  |  |  |  |  |  |  |  |
| insula | 18 | 29 | ** | ** | ** X |  |  |  |  |  |  |  |  |  |  |  |  |
| insula | 18 | 34 |  |  | ** |  |  |  |  | ** X |  |  |  |  |  |  |  |
| lateralorbitofrontal | 15 | 22 |  |  | ** X | ** |  | * | * |  |  | * |  |  |  |  |  |
| lateralorbitofrontal | 15 | 73 |  |  | * |  |  |  | ** |  |  |  |  |  |  | ** X |  |
| lateralorbitofrontal | 18 | 25 | * |  | * | * X |  |  |  | * |  |  |  |  |  |  |  |
| lateralorbitofrontal | 18 | 26 | ** | ** | ** | ** X |  |  |  | * |  |  |  |  |  |  |  |
| middletemporal | 11 | 45 |  | * | * X |  |  | * |  |  |  |  |  |  |  |  |  |
| middletemporal | 14 | 48 |  |  | * X |  |  |  |  |  |  |  |  |  |  |  |  |
| middletemporal | 16 | 24 |  |  | * |  |  |  | * |  |  |  |  | ** X |  |  |  |
| middletemporal | 18 | 134 |  |  | * | * X |  |  |  |  |  |  |  |  |  |  |  |
| parahippocampal | 6 | 8 |  |  | ** |  |  |  |  |  |  |  |  |  |  |  | ** X |
| parahippocampal | 6 | 9 |  |  | ** |  |  |  |  |  |  | ** |  |  |  |  | ** X |
| parahippocampal | 10 | 8 |  |  | * X |  |  |  |  |  |  |  |  |  |  |  |  |
| parsopercularis | 8 | 22 |  |  | ** X | ** |  |  | ** |  |  |  |  | * |  |  |  |
| parsopercularis | 8 | 23 |  |  | ** | ** X |  |  | ** |  |  |  |  |  |  | ** |  |
| parsopercularis | 11 | 78 |  |  | * | ** X |  | * |  |  |  |  |  | * |  |  | * |
| parsopercularis | 18 | 32 |  |  | * | ** X |  |  |  |  |  |  |  |  |  |  |  |
| parstriangularis | 10 | 59 |  |  | ** X |  |  |  |  |  |  |  |  |  |  |  |  |
| postcentral | 12 | 39 |  |  | * |  |  |  | * X |  |  | * |  |  |  |  |  |
| postcentral | 12 | 49 |  |  | ** X |  |  |  |  |  |  |  |  |  |  |  |  |
| precentral | 15 | 15 |  |  | * | ** X |  |  | ** |  |  |  |  |  |  |  |  |
| precentral | 15 | 16 |  |  | ** | ** X |  |  |  | ** |  |  |  |  |  |  |  |
| rostralmiddlefrontal | 8 | 24 |  |  | ** | ** X |  |  | ** |  |  |  |  |  |  | * |  |
| superiorfrontal | 12 | 6 |  |  | ** |  |  |  | ** X |  |  |  |  |  |  |  |  |
| superiorparietal | 3 | 111 |  |  | * |  |  |  | * |  | ** X |  |  |  |  |  |  |
| superiorparietal | 3 | 112 |  |  | * |  |  |  | * X |  |  |  |  |  |  |  |  |
| superiortemporal | 2 | 44 |  |  | * | ** X |  |  |  |  |  |  |  |  |  |  |  |
| supramarginal | 2 | 37 |  |  | * |  |  |  | ** X |  |  |  | ** |  |  |  |  |

**Table S4B.**
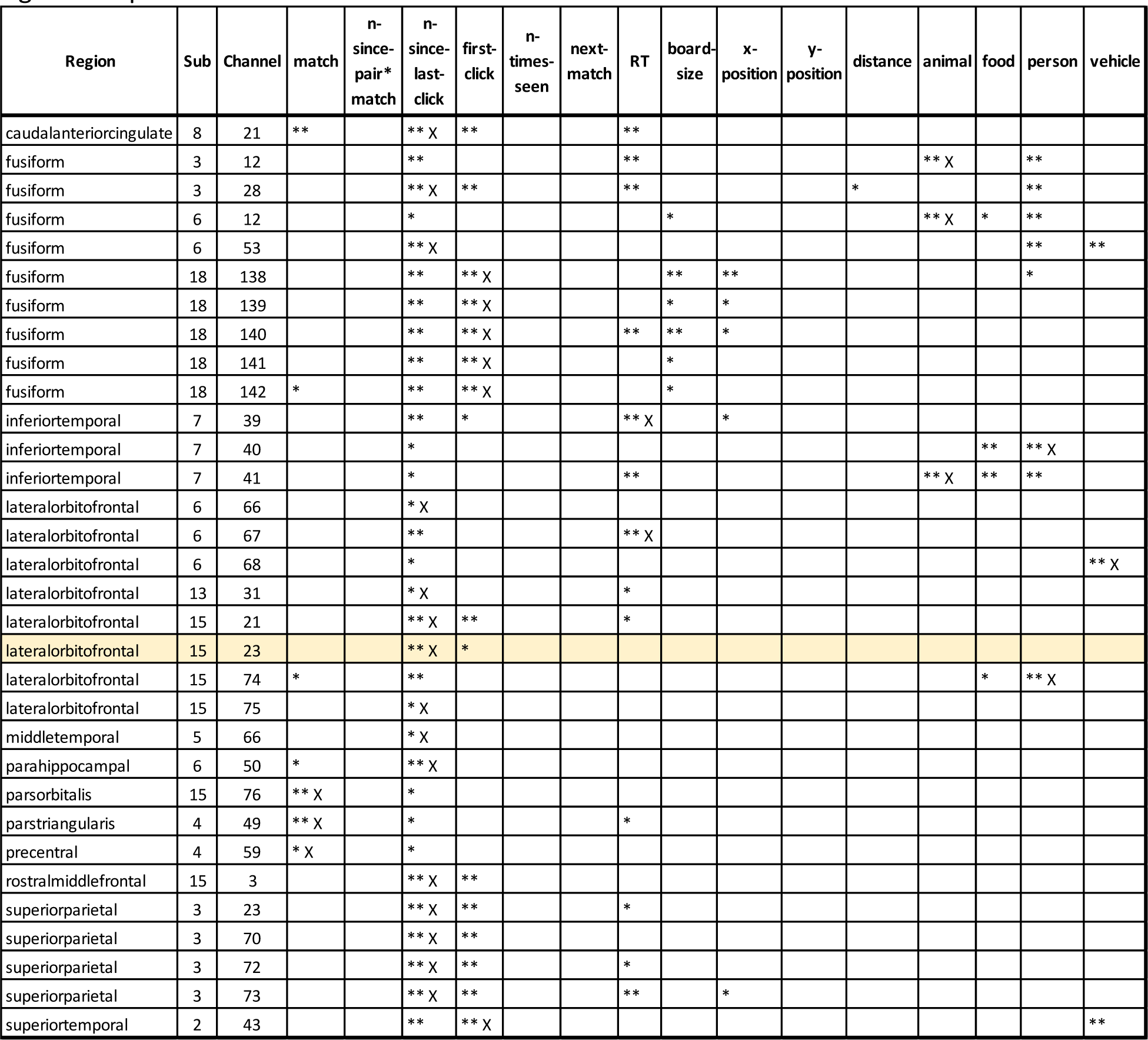
First tile GLM analysis for all white matter electrodes that showed n-since-last-click as a significant predictor. Each column (starting from column 4) shows a separate predictor. Yellow row indicates the example electrode in **FigureS4**. *: p<0.01. **: p<0.001. X: most significant predictor.

**Table S5A.**
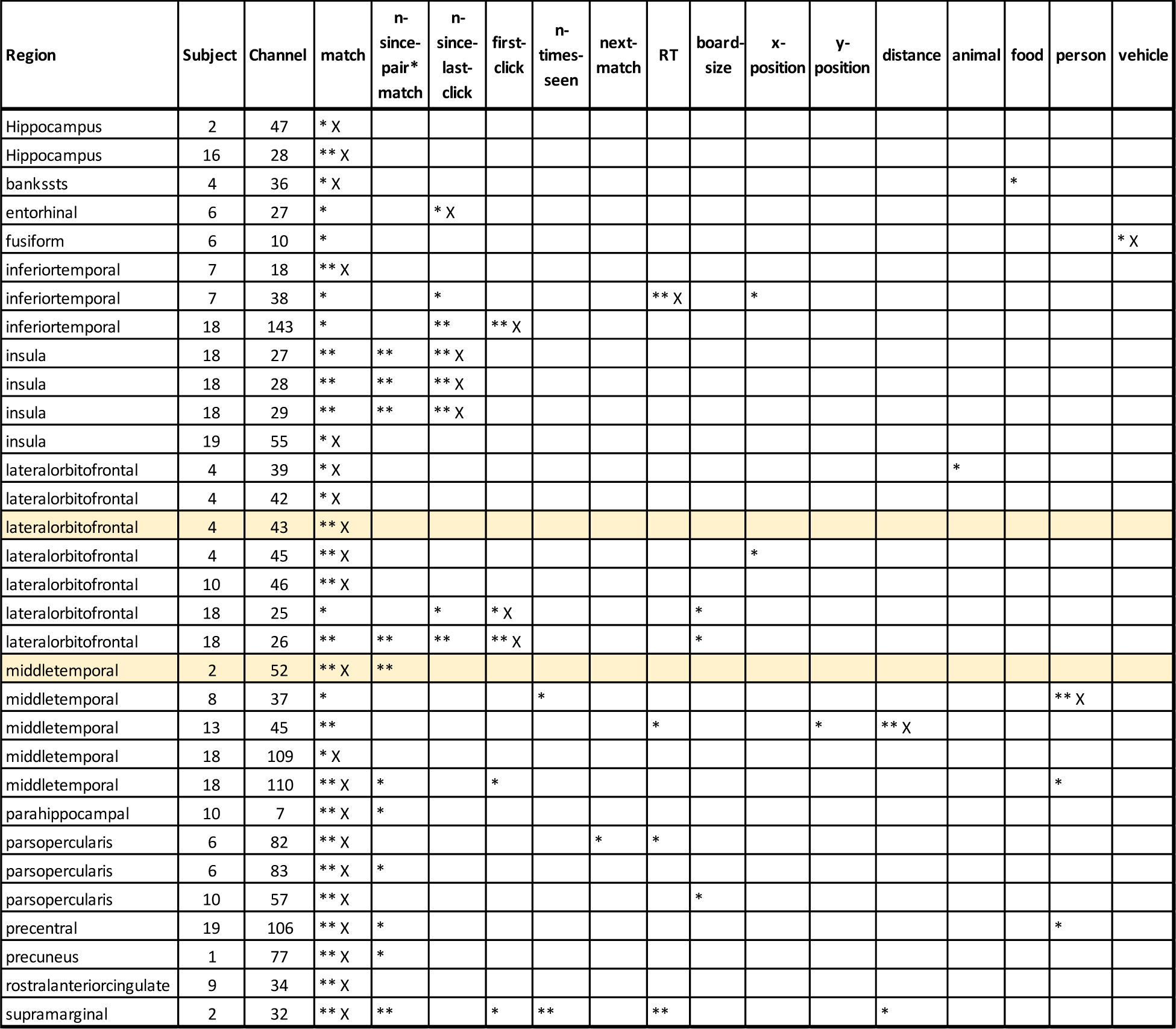
First tile GLM analysis for all gray matter electrodes that showed match as a significant predictor. Each column (starting from column 4) shows a separate predictor. Yellow rows indicate the example electrodes in **Figure 5** (first row), **Figure S7** (first row), and **Figure S8** (second row).*: p<0.01. **: p<0.001. X: most significant predictor.

**Table S5B.** First tile GLM analysis for all white matter electrodes that showed match as a significant predictor. Each column (starting from column 4) shows a separate predictor. Yellow row indicates the example electrode in **Figure S6**. *: p<0.01. **: p<0.001. X: most significant predictor.

| Region | Sub | Channel | match | n-since-pair*<br>match | n-since-last-click | first-click | n-times-seen | next-match | RT | board-size | x-position | y-position | distance | animal | food | person | vehicle |
| --- | --- | --- | --- | --- | --- | --- | --- | --- | --- | --- | --- | --- | --- | --- | --- | --- | --- |
| caudalanteriorcingulate | 8 | 21 | ** |  | ** X | ** |  |  | ** |  |  |  |  |  |  |  |  |
| entorhinal | 10 | 19 | ** X |  |  |  |  |  |  |  |  |  |  |  |  |  |  |
| fusiform | 18 | 142 | * |  | ** | ** X |  |  |  | * |  |  |  |  |  |  |  |
| inferiortemporal | 6 | 57 | ** X |  |  |  |  |  |  |  |  |  |  |  |  |  |  |
| inferiortemporal | 7 | 19 | * X |  |  |  |  |  |  |  |  |  |  |  |  |  |  |
| inferiortemporal | 7 | 63 | * X |  |  |  |  |  |  |  |  |  |  |  |  |  |  |
| lateralorbitofrontal | 4 | 41 | ** X |  |  |  |  |  |  |  |  |  |  |  |  |  |  |
| lateralorbitofrontal | 4 | 44 | * X |  |  |  |  |  |  |  |  |  |  |  |  |  |  |
| lateralorbitofrontal | 10 | 55 | ** X | ** |  |  | * |  |  |  |  |  |  |  |  |  |  |
| lateralorbitofrontal | 14 | 34 | ** X |  |  |  |  |  |  |  |  |  |  |  |  |  |  |
| lateralorbitofrontal | 14 | 36 | ** X | * |  |  | * |  |  |  |  |  |  |  |  |  | * |
| lateralorbitofrontal | 15 | 74 | * |  | ** |  |  |  |  |  |  |  |  |  | * | ** X |  |
| lingual | 11 | 60 | * |  |  |  |  |  |  |  | ** | * |  | ** |  | ** X |  |
| medialorbitofrontal | 11 | 63 | ** X |  |  |  |  | * |  |  |  |  |  |  |  |  |  |
| middletemporal | 9 | 24 | ** X |  |  |  |  |  |  |  |  |  |  |  |  |  |  |
| middletemporal | 13 | 44 | * |  |  |  |  |  | * |  |  | * | ** X |  |  |  |  |
| parahippocampal | 6 | 50 | * |  | ** X |  |  |  |  |  |  |  |  |  |  |  |  |
| parsopercularis | 6 | 81 | ** X | ** |  |  |  |  |  |  |  |  |  |  |  |  |  |
| parsorbitalis | 15 | 76 | ** X |  | * |  |  |  |  |  |  |  |  |  |  |  |  |
| parstriangularis | 4 | 49 | ** X |  | * |  |  |  | * |  |  |  |  |  |  |  |  |
| parstriangularis | 7 | 82 | ** X | * |  |  |  |  |  |  |  |  |  |  |  |  |  |
| parstriangularis | 7 | 84 | ** X | ** |  |  |  |  |  |  |  |  |  |  |  |  |  |
| precentral | 4 | 55 | ** X |  |  |  |  |  |  |  |  |  |  |  |  |  |  |
| precentral | 4 | 56 | ** X |  |  |  |  |  |  |  |  |  |  |  |  |  |  |
| precentral | 4 | 57 | ** X | * |  |  |  |  |  |  |  |  |  |  |  |  |  |
| precentral | 4 | 58 | ** X |  |  |  |  |  |  |  |  |  |  |  |  |  |  |
| precentral | 4 | 59 | * X |  | * |  |  |  |  |  |  |  |  |  |  |  |  |
| superiorparietal | 3 | 20 | * |  |  | * |  |  |  | * X |  |  |  |  |  |  |  |
| superiortemporal | 6 | 19 | * |  |  |  |  | * |  |  | * X |  | * |  |  |  |  |
| supramarginal | 2 | 31 | ** X | ** |  |  |  |  | * |  |  |  |  |  |  |  |  |

**Table S6A.** First tile GLM analysis for all gray matter electrodes that showed n-since-pair*match as a significant predictor. Each column (starting from column 4) shows a separate predictor. Yellow row indicates the example electrode in **Figure 6**.*: p<0.01. **: p<0.001. X: most significant predictor.

| Region | Sub | Channel | match | n-since-pair*match | n-since-last-click | first-click | n-times-seen | next-match | RT | board-size | x-position | y-position | distance | animal | food | person | vehicle |
| --- | --- | --- | --- | --- | --- | --- | --- | --- | --- | --- | --- | --- | --- | --- | --- | --- | --- |
| insula | 18 | 27 | ** | ** | ** X |  |  |  |  |  |  |  |  |  |  |  |  |
| insula | 18 | 28 | ** | ** | ** X |  |  |  |  |  |  |  |  |  |  |  |  |
| insula | 18 | 29 | ** | ** | ** X |  |  |  |  |  |  |  |  |  |  |  |  |
| lateralorbitofrontal | 18 | 3 |  | * X |  |  |  |  |  |  |  |  |  |  |  |  |  |
| lateralorbitofrontal | 18 | 6 |  | * |  | ** X |  |  |  | ** |  |  |  |  |  |  |  |
| lateralorbitofrontal | 18 | 26 | ** | ** | ** | ** X |  |  |  | * |  |  |  |  |  |  |  |
| medialorbitofrontal | 18 | 2 |  | * X |  |  |  |  |  |  |  |  |  |  |  |  |  |
| middletemporal | 2 | 52 | ** X | ** |  |  |  |  |  |  |  |  |  |  |  |  |  |
| middletemporal | 11 | 45 |  | * | * X |  |  | * |  |  |  |  |  |  |  |  |  |
| middletemporal | 18 | 110 | ** X | * |  | * |  |  |  |  |  |  |  |  |  | * |  |
| parahippocampal | 10 | 7 | ** X | * |  |  |  |  |  |  |  |  |  |  |  |  |  |
| parsopercularis | 6 | 83 | ** X | * |  |  |  |  |  |  |  |  |  |  |  |  |  |
| precentral | 19 | 106 | ** X | * |  |  |  |  |  |  |  |  |  |  |  | * |  |
| precuneus | 1 | 77 | ** X | * |  |  |  |  |  |  |  |  |  |  |  |  |  |
| supramarginal | 2 | 32 | ** X | ** |  | * | ** |  | ** |  |  |  | * |  |  |  |  |

**Table S6B.** First tile GLM analysis for all white matter electrodes that showed n-since- pair*match as a significant predictor. Each column (starting from column 4) shows a separate predictor. Yellow row indicates the example electrode in **Figure S9**. *: p<0.01. **: p<0.001. X: most significant predictor.

| Region | Sub | Channel | match | n-since-pair*match | n-since-last-click | first-click | n-times-seen | next-match | RT | board-size | x-position | y-position | distance | animal | food | person | vehicle |
| --- | --- | --- | --- | --- | --- | --- | --- | --- | --- | --- | --- | --- | --- | --- | --- | --- | --- |
| lateralorbitofrontal | 10 | 55 | ** X | ** |  |  | * |  |  |  |  |  |  |  |  |  |  |
| lateralorbitofrontal | 14 | 6 |  | ** X |  |  |  |  |  |  |  |  | * |  |  |  |  |
| lateralorbitofrontal | 14 | 36 | ** X | * |  |  | * |  |  |  |  |  |  |  |  |  | * |
| middletemporal | 18 | 60 |  | * |  |  |  |  |  | * X |  | * |  |  |  |  |  |
| parsopercularis | 6 | 81 | ** X | ** |  |  |  |  |  |  |  |  |  |  |  |  |  |
| parstriangularis | 7 | 82 | ** X | * |  |  |  |  |  |  |  |  |  |  |  |  |  |
| parstriangularis | 7 | 84 | ** X | ** |  |  |  |  |  |  |  |  |  |  |  |  |  |
| precentral | 4 | 57 | ** X | * |  |  |  |  |  |  |  |  |  |  |  |  |  |
| supramarginal | 2 | 31 | ** X | ** |  |  |  |  | * |  |  |  |  |  |  |  |  |

**Table S7A.** Second tile GLM analysis for all gray matter electrodes that showed first-click as a significant predictor. Each column (starting from column 4) shows a separate predictor. *: p<0.01. **: p<0.001. X: most significant predictor.

| Region | Sub | Channel | match | n-since-last-click | first-click | n-times-seen | next-match | RT | board-size | x-position | y-position | distance | animal | food | person | vehicle |
| --- | --- | --- | --- | --- | --- | --- | --- | --- | --- | --- | --- | --- | --- | --- | --- | --- |
| Amygdala | 10 | 10 |  |  | * X |  |  |  |  |  |  |  |  |  |  |  |
| Hippocampus | 18 | 104 | ** X |  | ** |  |  |  |  |  |  |  |  |  |  |  |
| bankssts | 18 | 131 | * |  | * |  |  | ** X | * |  |  |  |  |  |  |  |
| fusiform | 3 | 11 |  | * | * |  |  |  |  |  |  |  | ** X |  |  |  |
| fusiform | 10 | 37 |  |  | * |  |  |  |  |  |  |  |  |  |  | ** X |
| insula | 18 | 27 | ** X |  | ** | * |  | * |  |  |  |  |  |  |  |  |
| insula | 18 | 28 | ** X |  | * | * |  |  |  |  |  |  |  |  |  |  |
| lateraloccipital | 20 | 4 |  |  | * X |  |  |  |  |  |  |  |  |  |  |  |
| lateraloccipital | 20 | 5 |  |  | ** X |  |  |  |  |  |  |  |  |  |  |  |
| lateralorbitofrontal | 15 | 22 | * | ** | ** |  | * |  |  |  |  |  |  | ** |  | ** X |
| lateralorbitofrontal | 18 | 5 |  |  | * X |  |  |  | * |  |  |  |  |  |  |  |
| lateralorbitofrontal | 18 | 25 | ** |  | ** X | ** |  | ** |  |  |  |  |  |  |  |  |
| lateralorbitofrontal | 18 | 26 | ** | * | ** X | ** |  | ** |  |  |  |  |  |  |  |  |
| lingual | 1 | 12 | * |  | * |  |  |  |  |  | ** X |  |  | * |  |  |
| parahippocampal | 9 | 27 |  |  | * |  |  |  | ** X |  |  |  |  |  |  |  |
| parsopercularis | 8 | 22 | ** | ** | ** |  |  | ** X | ** |  |  |  | ** |  |  |  |
| parsopercularis | 8 | 23 | ** |  | ** |  |  | ** X | ** |  |  |  |  |  |  |  |
| postcentral | 20 | 83 |  |  | * X |  |  |  |  | * |  |  |  |  |  |  |
| postcentral | 20 | 84 |  | * | * X |  |  |  |  |  |  |  |  |  |  |  |
| precentral | 18 | 40 |  |  | * X | * |  |  |  |  |  | * |  |  |  |  |
| precentral | 12 | 24 | * | * | * X |  |  |  |  |  |  |  |  |  |  |  |
| precuneus | 1 | 77 |  |  | * |  | ** X |  |  |  |  |  |  |  |  |  |
| rostralmiddlefrontal | 8 | 24 | ** X | * | ** |  |  | ** |  |  |  |  |  |  |  |  |
| superiortemporal | 18 | 119 |  |  | * X |  |  |  |  |  | * |  |  |  |  |  |

**Table S7B.** Second tile GLM analysis for all white matter electrodes that showed first-click as a significant predictor. Each column (starting from column 4) shows a separate predictor. *: p<0.01. **: p<0.001. X: most significant predictor.

| Region | Sub | Channel | match | n-since-last-click | first-click | n-times-seen | next-match | RT | board-size | x-position | y-position | distance | animal | food | person | vehicle |
| --- | --- | --- | --- | --- | --- | --- | --- | --- | --- | --- | --- | --- | --- | --- | --- | --- |
| caudalanteriorcingulate | 8 | 21 | ** | ** | * |  |  | ** X |  |  |  |  |  |  |  |  |
| fusiform | 18 | 138 | ** X | ** | * |  |  |  | ** | * |  |  |  |  | * |  |
| inferiorparietal | 20 | 15 |  |  | ** X |  |  |  |  |  |  |  |  |  |  |  |
| lateralorbitofrontal | 6 | 68 |  | * | ** X | * |  |  |  |  |  |  |  |  |  |  |
| lateralorbitofrontal | 15 | 23 |  | * | ** |  |  |  |  |  |  |  |  |  |  | ** X |
| lateralorbitofrontal | 18 | 24 | ** | * | ** X | * |  |  |  |  |  |  |  |  |  |  |
| parstriangularis | 16 | 7 |  |  | ** X |  |  |  |  |  |  |  |  |  |  |  |
| parstriangularis | 16 | 9 |  |  | ** X |  |  |  |  |  |  |  |  |  |  |  |
| posteriorcingulate | 20 | 123 |  |  | ** X |  |  | * |  |  |  |  |  |  |  |  |
| precentral | 4 | 54 |  |  | * X |  |  |  |  |  |  |  |  |  |  |  |
| precentral | 4 | 57 |  |  | * |  | * X |  |  |  |  |  |  |  |  |  |
| precentral | 4 | 58 |  |  | * |  | ** X |  |  |  |  |  |  |  |  |  |

**Table S8A.** Second tile GLM analysis for all gray matter electrodes that showed n-since-last-click as a significant predictor. Each column (starting from column 4) shows a separate predictor. Yellow row indicates the example electrode in **Figure S10**. *: p<0.01. **: p<0.001. X: most significant predictor.

| Region | Sub | Channel | match | n-since-last-click | first-click | n-times-seen | next-match | RT | board-size | x-position | y-position | distance | animal | food | person | vehicle |
| --- | --- | --- | --- | --- | --- | --- | --- | --- | --- | --- | --- | --- | --- | --- | --- | --- |
| fusiform | 3 | 11 |  | * | * |  |  |  |  |  |  |  | ** X |  |  |  |
| lateralorbitofrontal | 122 | 46 |  | * X |  |  |  |  |  |  |  |  |  |  |  |  |
| lateralorbitofrontal | 15 | 22 | * | ** | ** |  | * |  |  |  |  |  |  | ** |  | ** X |
| lateralorbitofrontal | 18 | 9 | ** X | * |  |  |  | ** | * |  |  |  |  |  |  |  |
| lateralorbitofrontal | 18 | 26 | ** | * | ** X | ** |  | ** |  |  |  |  |  |  |  |  |
| parsopercularis | 8 | 22 | ** | ** | ** |  |  | ** X | ** |  |  |  | ** |  |  |  |
| postcentral | 20 | 84 |  | * | * X |  |  |  |  |  |  |  |  |  |  |  |
| precentral | 12 | 24 | * | * | * X |  |  |  |  |  |  |  |  |  |  |  |
| rostralmiddlefrontal | 8 | 24 | ** X | * | ** |  |  | ** |  |  |  |  |  |  |  |  |

**Table S8B.** Second tile GLM analysis for all white matter electrodes that showed n-since-last-click as a significant predictor. Each column (starting from column 4) shows a separate predictor. *: p<0.01. **: p<0.001. X: most significant predictor.

| Region | Sub | Channel | match | n-since-last-click | first-click | n-times-seen | next-match | RT | board-size | x-position | y-position | distance | animal | food | person | vehicle |
| --- | --- | --- | --- | --- | --- | --- | --- | --- | --- | --- | --- | --- | --- | --- | --- | --- |
| caudalanteriorcingulate | 8 | 21 | ** | ** | * |  |  | ** X |  |  |  |  |  |  |  |  |
| caudalmiddlefrontal | 3 | 90 |  | ** X |  |  | * |  |  |  |  |  |  |  |  |  |
| fusiform | 3 | 28 |  | ** |  |  |  | * |  |  |  |  |  |  | ** X |  |
| fusiform | 6 | 12 | ** | ** |  |  |  |  |  |  |  |  | ** X | * | * |  |
| fusiform | 6 | 53 |  | * |  |  |  | ** |  |  |  |  |  |  | ** X | * |
| fusiform | 18 | 138 | ** X | ** | * |  |  |  | ** | * |  |  |  |  | * |  |
| fusiform | 18 | 139 | ** X | ** |  |  |  | * |  | * |  |  |  |  |  |  |
| fusiform | 18 | 140 | ** X | * |  |  |  | ** |  | ** |  |  |  |  |  |  |
| fusiform | 18 | 141 | ** X | ** |  |  |  | ** |  | * |  |  |  |  |  |  |
| lateralorbitofrontal | 6 | 68 |  | * | ** X | * |  |  |  |  |  |  |  |  |  |  |
| lateralorbitofrontal | 15 | 21 | ** X | * |  |  |  |  |  |  |  |  |  |  |  |  |
| lateralorbitofrontal | 15 | 23 |  | * | ** |  |  |  |  |  |  |  |  |  |  | ** X |
| lateralorbitofrontal | 15 | 74 |  | * |  |  |  | ** X |  |  |  |  |  |  | * |  |
| lateralorbitofrontal | 18 | 24 | ** | * | ** X | * |  |  |  |  |  |  |  |  |  |  |
| middletemporal | 7 | 10 |  | * X |  |  |  |  |  |  |  |  |  |  |  |  |
| middletemporal | 7 | 66 |  | * |  |  |  |  |  |  |  |  |  | * X | * |  |
| middletemporal | 7 | 67 | * X | * |  |  |  |  |  |  | * |  |  |  |  |  |
| parahippocampal | 6 | 50 | * | ** X |  |  |  |  |  |  |  |  |  |  |  |  |
| parsorbitalis | 6 | 71 |  | * X |  |  |  |  |  |  |  |  |  |  |  |  |
| parsorbitalis | 15 | 76 |  | * |  |  |  | * |  |  |  |  |  |  |  | ** X |
| parstriangularis | 2 | 28 |  | * X |  |  |  |  |  |  |  |  |  |  |  |  |
| parstriangularis | 2 | 29 |  | * |  |  |  |  |  |  |  |  |  |  |  | * X |
| parstriangularis | 7 | 83 | ** X | * |  |  | * |  |  |  |  |  |  |  |  |  |
| superiorparietal | 3 | 72 | ** X | * |  |  |  |  | * |  |  |  |  |  | * |  |

**Table S9A.**
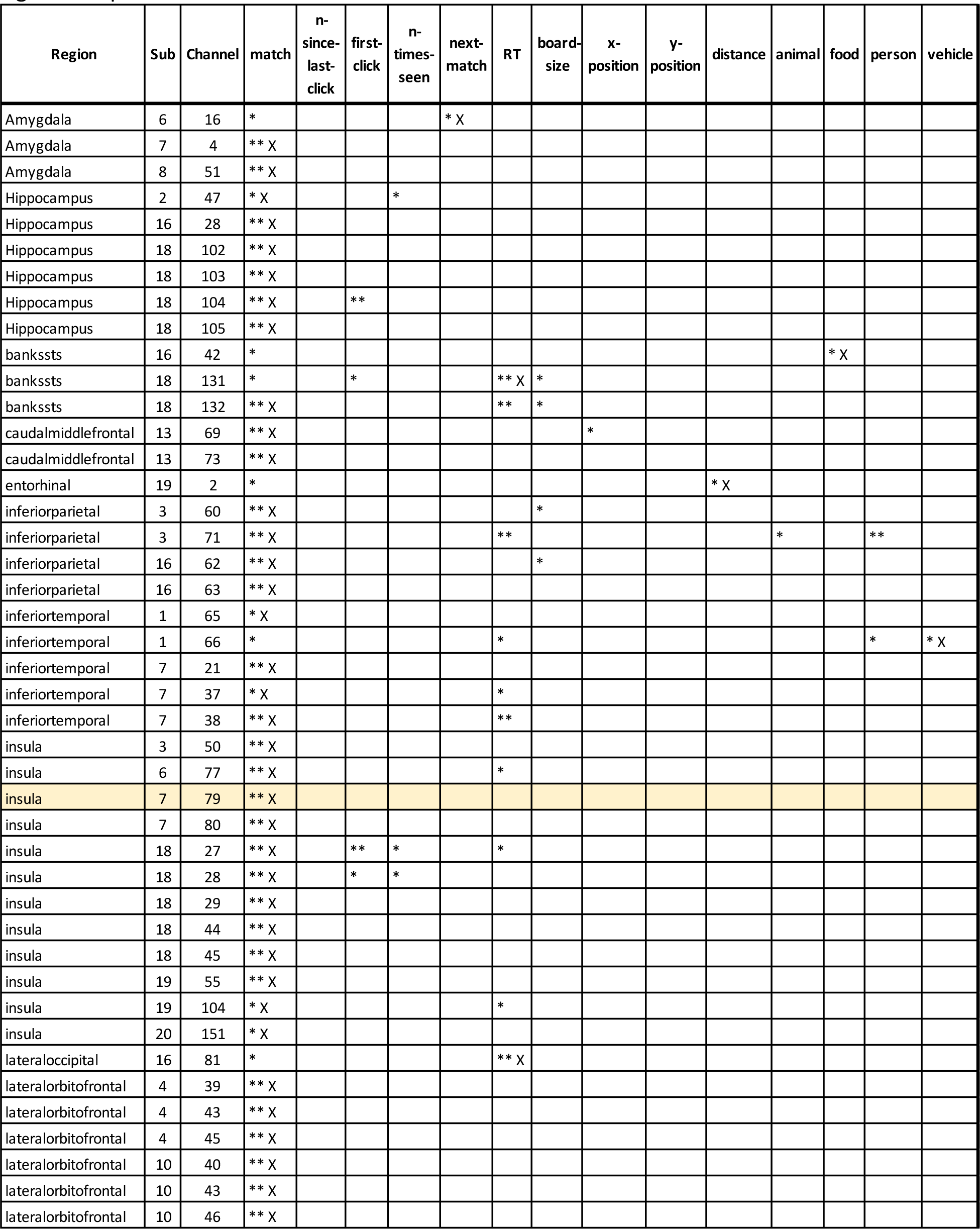
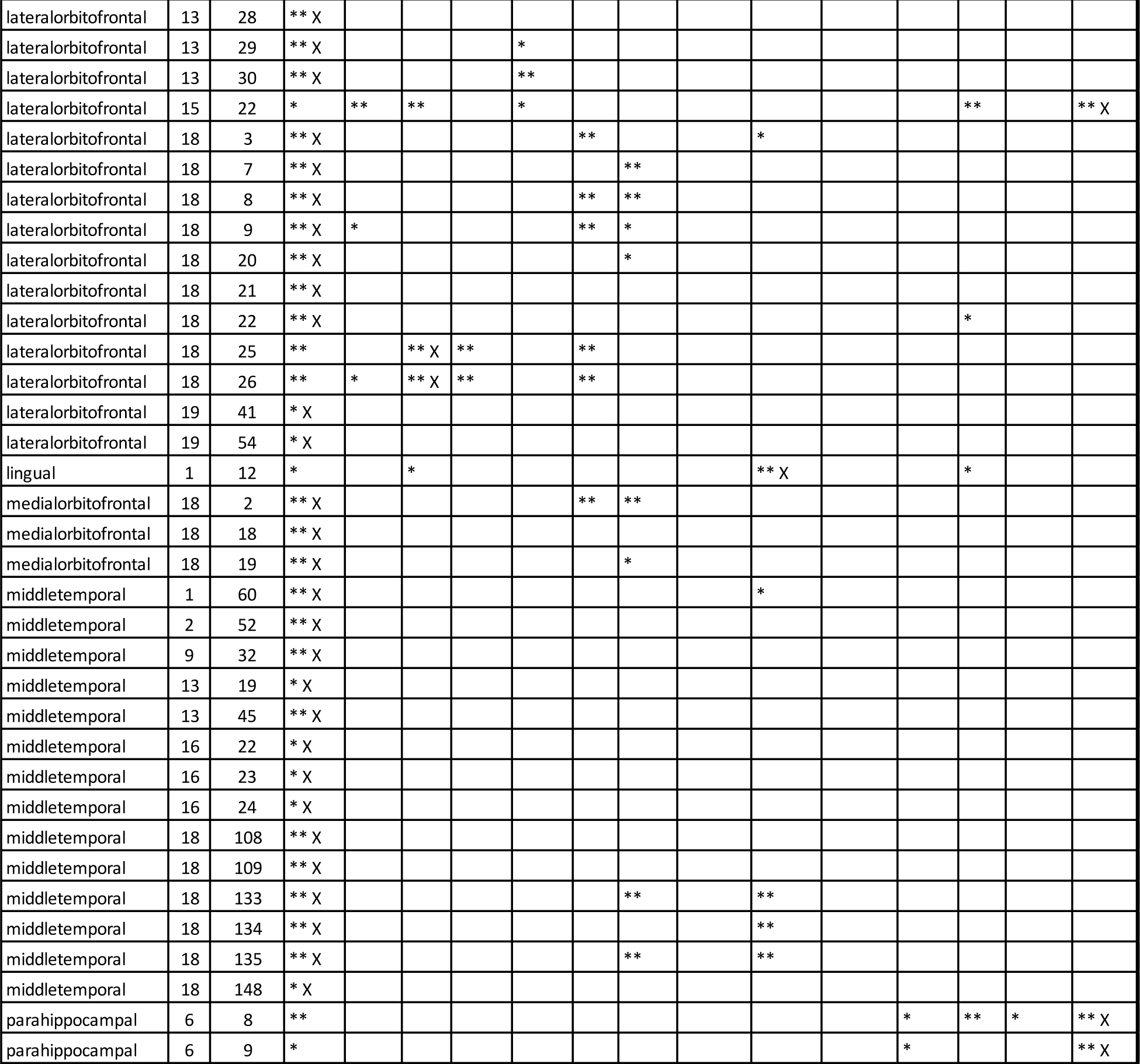
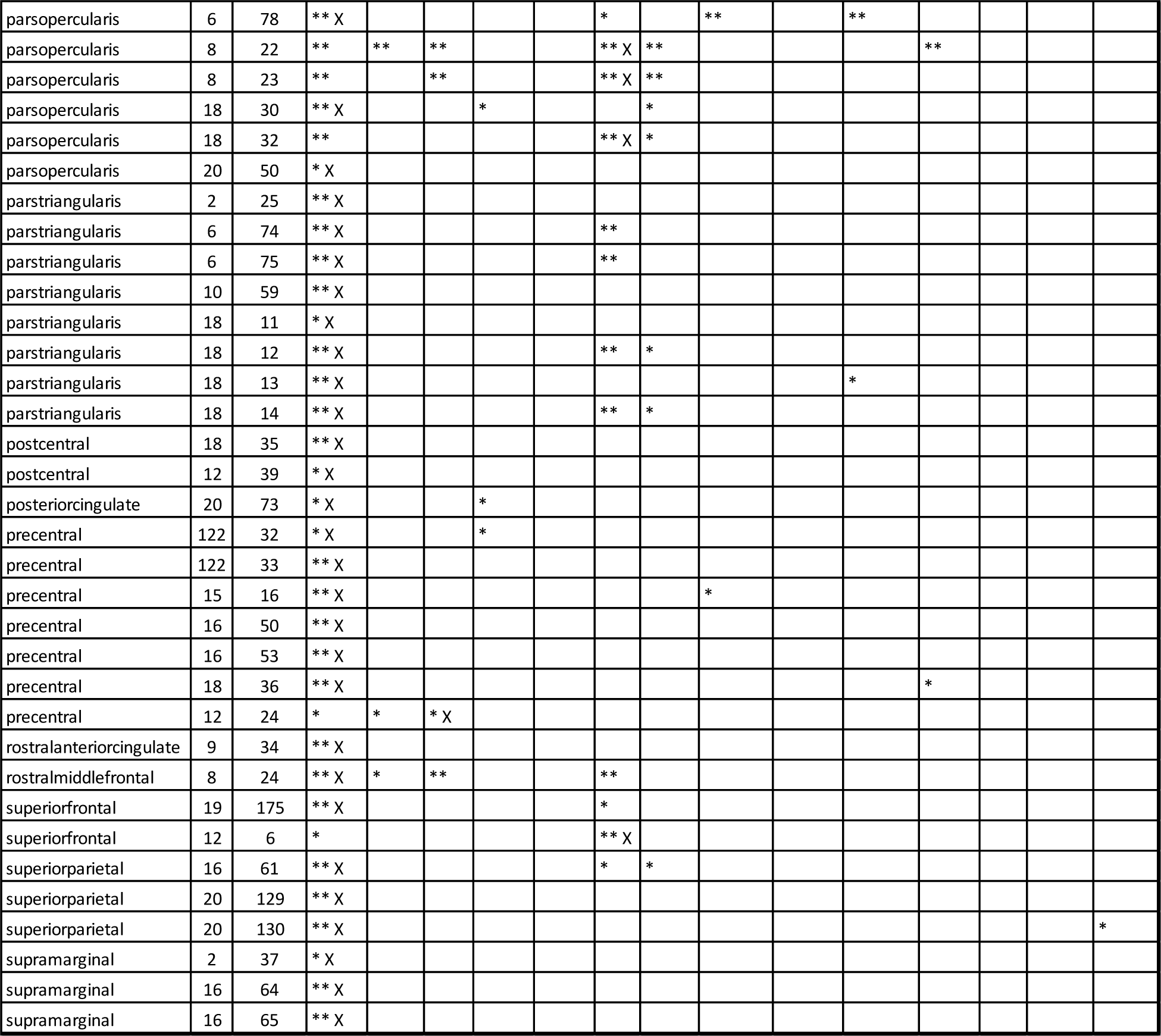
Second tile GLM analysis for all gray matter electrodes that showed match as a significant predictor. Each column (starting from column 4) shows a separate predictor. Yellow row indicates the example electrode in **Figure 7** and **Figure S9**. *: p<0.01. **: p<0.001. X: most significant predictor.

**Table S9B.**
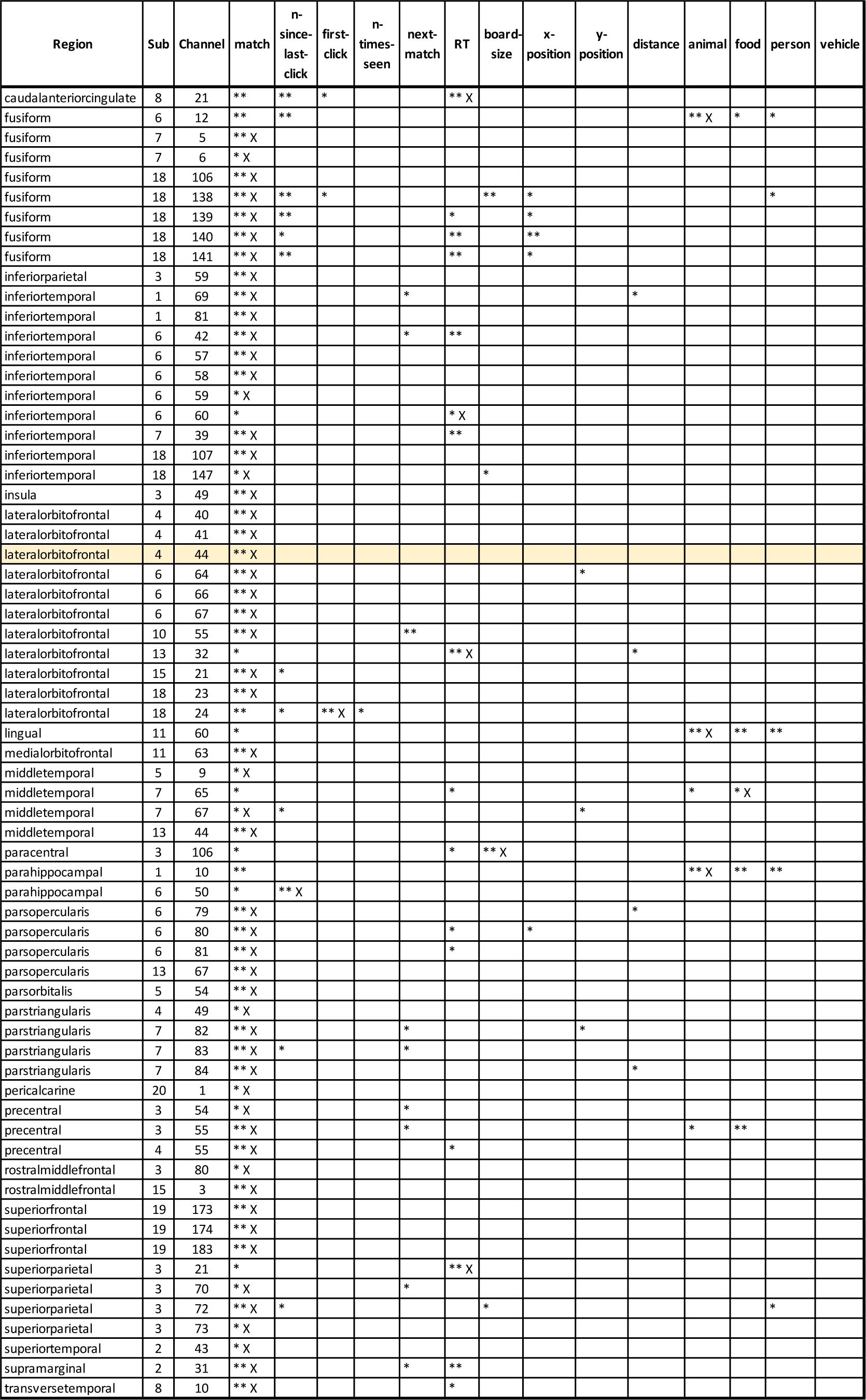
Second tile GLM analysis for all white matter electrodes that showed match as a significant predictor. Each column (starting from column 4) shows a separate predictor. Yellow row indicates the example electrode in **Figure S11**. *: p<0.01. **: p<0.001. X: most significant predictor.

## Supplementary Video 1

This video shows the sequence of events in the task for boards of different size for one of the participants in the experiment. In each trial, participants click on two tiles. If they match, the tiles turn green, otherwise, the tiles turn black again. The goal is to find all matching pairs. See **Methods** for an expanded description of the task.

